# Compensatory Mechanisms in γδ T Cell-Deficient Chickens Following *Salmonella* infection

**DOI:** 10.1101/2025.02.11.637636

**Authors:** Felix Tetzlaff, Ulrich Methner, Theresa von Heyl, Christian Menge, Benjamin Schusser, Angela Berndt

## Abstract

Avian γδ T lymphocytes are highly abundant in the intestinal mucosa and play a critical role in immune defense against infectious diseases in chickens. However, their specific contributions to infection control remain poorly understood. To investigate the role of γδ T cells and their possible compensation, we studied wild-type and γδ T cell knockout chickens following infection with *Salmonella* Enteritidis. Bacterial loads in the liver, cecal content, and cecal wall were quantified. Immune cell populations in blood, spleen, and cecum were analyzed using flow cytometry. Immune gene transcription in sorted T cell subsets and cecal tissue was measured by RT-qPCR.

Strikingly, chickens lacking γδ T cells had significantly higher bacterial loads in the liver and more extensive *Salmonella* invasion in the cecal wall during the early stages of infection compared to wild-type birds. In the blood, infected γδ T cell knockout chickens displayed a significantly increased percentage of CD25^+^ NK-like cells. In both blood and tissue, infected wild-type chickens demonstrated an increased absolute number of CD8αα^++^ γδ T cells. Conversely, γδ T cell knockout chickens exhibited an augmented cell count of a CD8αα^++^CD4^-^ non-γδ T cell population after infection, which might include αβ T cells. At 7 days post infection (dpi), gene expression analysis revealed elevated transcription of the activation marker IL-2Rα and proinflammatory cytokines (IL-17A, IFN-γ) in CD8αα^++^CD4^-^ non-γδ T cells from γδ T cell knockout chickens compared to CD8αα^++^ γδ T cells from wild-type birds. By 12 dpi, these differences diminished as transcription levels increased in γδ T cells of wild-type animals.

Our findings demonstrate that γδ T cells play a role in early immune protection against *Salmonella* Enteritidis infection in chickens. In later stages of the infection, the γδ T cells and their functions appear to be replaced by other cells.

## 1. Introduction

*Salmonella* (*S*.) spp. is a predominant cause of foodborne gastrointestinal infection within the European Union, with 65,208 confirmed cases of human salmonellosis reported in 2022 (1). Notably, *S.* Enteritidis was identified as prominent pathogen, accounting for around 67% of reported cases and posing a global health concern. The main sources are poultry-derived products, mainly eggs and egg products but also chicken meat (1). For this reason, the primary aim of *Salmonella* control is to prevent these organisms from entering the food chain. This is achieved by stringent hygiene regimes, novel vaccine programs and selection of chicken breeds with enhanced colonization resistance. However, susceptibility to *S.* infection varies with age in poultry, with chicks younger than 3 days showing increased vulnerability due to their not yet fully developed gut microbiota and immune system (2–5). This underlines the crucial need for in-depth research into the avian immune response to *Salmonella* (*S*.) infection for improving chicken protection and minimizing human exposure to the pathogen in the future.

*S.* Enteritidis is now as ever highly prevalent in poultry, especially in laying hens. This serovar is able to induce both extensive intestinal colonization as well as pronounced invasion and potential systemic spread to different organs including the reproductive tract. The avian immune response in cecum, spleen, bursa of Fabricius and blood is marked by a significant influx of immune cells. Heterophils act as initial effector response, whereas macrophages modulate innate immunity and facilitate development of adaptive immune mechanisms (6–11). T lymphocytes are key players in mediating immune responses against a broad spectrum of pathogens. Common to their mammalian counterparts, avian T cells exhibit heterodimeric T cell receptors (TCR), which consist of a constant and variable region with either alpha and beta chains (αβ T cells) or gamma and delta chains (γδ T cells) (12, 13). The functionality of αβ T cells, defined by the two co-receptors CD4 and CD8, is well-documented in many species, also in chickens (14–16). Avian CD8^+^ T cells are split into two subsets, characterized by surface expression of either CD8αα homo- or CD8αβ heterodimers.

In chickens, γδ T cells represent up to 50% of the peripheral T cell repertoire (17, 18), while in humans, mouse and rats, the frequency of γδ T cells is relatively low, ranging from 1 to 10% (19). Different species-specific subpopulations of γδ T cells have been described, depending on their localization, antigen expression and cytokine production. In chickens, γδ T cells (TCR1^+^ cells) are categorized into CD8α^++^ (high), CD8α^+^ (dim), and CD8α^-^ (negative) populations, displaying substantial subset and functional heterogeneity (20).

The γδ T cells are of special importance at mucosal sites of the body (21–23). They stimulate early immune defense mechanisms (24, 25) and may serve as a bridge between the innate and adaptive reactions (26). The γδ T cells are crucial for immune regulation, tissue healing, inflammation, and protection against intra- and extracellular pathogens. Moreover, γδ T cells have memory abilities and produce a variety of cytokines (27–32).

Despite the high frequency of γδ T cells in chickens, our understanding of their functions is limited. Studies have shown a rapid expansion of γδ T cells in response to bacterial and viral infections (33, 34). Avian γδ T cells can produce cytokines and interferons such as IL-2, IL-17, IL-10, and IFN-γ (34–36), and demonstrate cytotoxic activities (37).

In *Salmonella*-infected chickens, distinct CD8α-expressing γδ T cell subsets quickly appear at inflammatory sites (9, 38). Especially the percentage of CD8αα^++^ (high) γδ T cells expand (34, 38). These cells express specific activation markers including CD25 and can produce immune effectors, such as IFN-γ and IL-2 (20, 39). However, whether γδ T cells counteract transepithelial invasion and dissemination of *Salmonella* and whether γδ T cell subsets exert different functions during the immune defense against *Salmonella* remains elusive in chickens. To study the immunological role of γδ T cells, a chicken line lacking γδ T cells have been developed. Complete absence of γδ T cells was achieved through genomic deletion of the constant region of the T cell receptor γ (TCR Cy^-/-^). The γδ T cell knockout chickens show no pathological phenotype and are indistinguishable from wild-type birds in both behavior and appearance (40).

The absence of γδ T cells can have divergent consequences at different stages of the host immune response. Mice with γδ T cell deficiencies (due to gene targeting or antibody-mediated depletion) exhibit inadequate early innate resistance to infections such as *L. monocytogenes* and

*C. albicans*. This has been attributed to the reduced production of IFN-γ and IL-17A and a decreased recruitment of innate immune cells (41–44). Other authors report an exaggerated inflammatory response after *L. monocytogenes* infection of γδ T cell-deficient mice, probably mediated by reduced regulation of macrophage homeostasis, T cell activity and reduced IL-10 production (28, 45–47). Remarkably, in uninfected mice, other immune cells, such as αβ T cells, appear to occupy the niches left by γδ T cells and partially compensate for their functions (48, 49).

To date, there have been no studies to determine whether and to what extent γδ T cells have a decisive influence on the course of the infection and the recovery of chickens after *Salmonella* exposure. In addition, it remains to be clarified whether the immunological functions of γδ T cells can somehow be compensated for in *Salmonella*-infected γδ T cell knockout chickens. Studies on γδ T cell function after *Salmonella* infection of γδ T cell knockout mice have produced conflicting results. Weintraub et al. (50) reported that intraepithelial γδ T cells (γδ IELs) and other intestinal γδ T cells do not play a significant role in controlling *Salmonella* invasion or replication in the mouse intestine. Conversely, Edelblum et al. (51) demonstrated that mice lacking γδ IELs exhibited increased *Salmonella* invasion and severity of systemic salmonellosis.

In this study, we set to address the role of γδ T cells in protecting chickens against the *Salmonella* infection by utilizing TCR Cy^-/-^ chickens lacking γδ T cells and wild-type chickens. Information on the dynamics of *Salmonella* dissemination in infected γδ T cell knockout and wild-type animals, alongside comparative analyses of the immunological responses in blood and tissues should be gained, contributing to a better understanding of γδ T cells and their possible redundancy in avian immune defense. We found a significance of γδ T cells in the course of the *Salmonella* infection in young chickens, and identified compensatory mechanisms in knockout chickens that maintain immune protection post infection by partially replacing functional properties of γδ T cells.

## 2. Materials and methods

### 2.1 Chickens

White Leghorn chickens with different genetic modifications were used for the experiments: Chickens with a homozygous knockout of the constant region of the T cell receptor-γ chain (TCR Cγ^-/-^, γδ T cell knockout chickens; (40)) and their non-modified hatch-siblings (wild-type chickens). The embryonated chicken eggs were produced at the TUM Animal Research Center, and hatched at the facilities of the Friedrich-Loeffler-Institut Jena. Genotyping was performed as described before (40). The animals were kept in cages. To effectively prevent cross-contamination between the groups, the individual groups were reared and kept in separate negative pressure rooms under standardized conditions and in accordance with the European Community Guidelines for Animal Welfare throughout the experiments. Antibiotic-free commercial feed in powder and pellet form and drinking water were available ad libitum. The study was conducted in strict compliance with the German Animal Welfare Act. The protocol was approved by the Ethics Committee for Animal Experiments and Animal Welfare of the State of Thuringia, Germany (registration number: BFI-20-001). Furthermore, the study was conducted under the supervision of the institutional animal welfare officer.

### 2.2 Bacteria

Oral infection of the wild-type and γδ T cell knockout animals was carried out with a rifampicin

(R) resistant variant of the comprehensively characterized strain *Salmonella* (*S.*) *enterica* subspecies *enterica* serovar Enteritidis 147 (SE147, phage type 4). The potential of this strain to colonize the cecum and invade internal organs of chickens has been reported in detail (9, 52–54). The strain had been stored in a cryobank system (Mast Diagnostica, Reinfeld, Germany) at −20 °C. The infection doses were estimated by measuring the absorbance at 600 nm using a calibration graph and subsequently confirmed by plate counting on nutrient agar (SIFIN, Berlin, Germany).

### 2.3 Experimental design

Two animal experiments were conducted (A, B). In each experiment, two groups of chickens were included, one group of wild-type and one group of γδ T cell knockout chickens. In experiment A, the chickens (64 wild-type and 37 TCR Cγ^-/-^ animals) were orally infected with *S.* Enteritidis 147 R at a dose of 2 x 10^7^ cfu/bird on day 3 of age. Oral administration was performed by instillation into the birds’ crop using syringes with an attached flexible tube. The volume of bacterial suspension used was 0.1 mL per bird. Three-day-old chickens of the control group (experiment B; 58 wild-type and 45 TCR Cγ^-/-^ animals) were mock-infected with 0.1 mL phosphate-buffered saline (PBS).

At eight distinct time points post infection, 1-13 birds per group were sacrificed and dissected. From each animal, the cecal content and tissue samples of spleen, liver, cecum and peripheral blood (EDTA coated Monovette®, Sarstedt, Nümbrecht, Germany) were taken. Samples were either analyzed immediately (cecal content, cecal tissue, liver, blood, spleen) or frozen and stored in liquid nitrogen or RNA-later (Qiagen, Hilden, Germany) for subsequent analysis (cecal tissue).

### 2.4 Bacteriology

Bacterial counts of *S*. Enteritidis 147 R were determined in cecal contents and in liver of all birds at each time point by a standard plating method (53–56). Briefly, homogenized liver tissue and cecal content were diluted in PBS, plated on deoxycholate-citrate agar (SIFIN) supplemented with rifampicin (100 µg/mL) and incubated at 37 °C for 18-24 h to detect bacterial colonization. Liver tissue and cecal content were also pre-enriched in buffered peptone water (SIFIN), incubated at 37 °C for 18-24 h and streaked onto deoxycholate-citrate agar supplemented with rifampicin (SIFIN).

### 2.5 Immune cell isolation

Immune cells were isolated from peripheral blood (PBMCs) for cell sorting and analysis of natural killer-like cells (NK-like cells) as well as from spleen and cecum for determination of absolute (leukocytes and T cell subsets) and relative (NK-like cells) immune cell numbers by flow cytometry.

To obtain the PBMCs, 1 mL EDTA-blood was mixed with 1 mL PBS plus 1% bovine serum albumin (BSA: Bio&Sell, Feucht, Germany; PBS^+^) and transferred on Pancoll animal separation medium (1.077 g/mL; Pan-Biotech, Aidenbach, Germany; Pancoll), followed by centrifugation at 800 x g for 20 min. The cell layer at the interface between PBS^+^ and Pancoll was transferred to a new centrifuge tube and washed with 5 mL PBS^+^ at 400 x g for 15 min. After removal of the supernatant, the cells were adjusted to 5 x 10^6^ cells/mL in PBS^+^ for subsequent flow cytometric staining and cell sorting.

To isolate splenic cells, the spleen was first weighed. The tissue was then placed in a petri dish containing 1 mL of sterile PBS^+^ and minced into small pieces using scalpel and forceps. Excess connective tissue was discarded. The remaining cell suspension was homogenized by slowly pipetting up and down using a 23G cannula (BD Bioscience, Heidelberg, Germany) with a syringe (B. Braun SE, Melsungen, Germany). After adding 1 mL fresh PBS^+^, the cell suspension was overlaid on Pancoll (1.077 g/mL) in a 12 mL tube and centrifuged at 800 x g for 20 min. The cell layer was transferred to a new centrifuge tube and washed with 5 mL PBS^+^ at 400 x g for 10 min. After removal of the supernatant, the cells were resuspended in 1 mL PBS^+^ and further diluted if necessary (to 5 x 10^6^ cells/mL) for subsequent flow cytometric staining.

To obtain the immune cells from the cecum, the tubular cecum was cut open, spread out for area measurement and washed thoroughly in PBS^+^ to remove the remaining cecal content. Epithelial and lamina propria cells were carefully scraped from the remaining muscular and serosa layers, the latter were subsequently removed. One mL of PBS^+^ was added, and the cell material was mechanically shredded with a scalpel. Next, 1 mL of collagenase type V solution from *Clostridium histolyticum* (2 mg/mL; Merck, Darmstadt, Germany) was added to the cells and incubated for 30 min at 37 °C on a shaker. The cells were then gently pipetted three times through a 1 mL pipette tip, transferred on 2 mL of Pancoll (1.077 g/mL; Pan-Biotech) and centrifuged at 800 x g for 20 min. The cell layer was transferred to a new tube and washed twice with 3 mL of fresh PBS^+^ at 400 x g for 10 min. Finally, the cell pellet was resuspended in 1 mL PBS^+^ and further diluted if necessary (to 5 x 10^6^ cells/mL) for subsequent flow cytometric staining.

### 2.6 Flow cytometry

To assess immune cell numbers in whole blood, PBMCs, spleen and cecum, we conducted flow cytometric analyses as previously described (57–59).

Four distinct antibody panels were employed for immune cell characterization (Table 1). Antibody mix 1 was designed to detect absolute numbers of leukocytes (thrombocytes, monocytes, T cells, B cells, granulocytes) of infected and mock-infected animals (58). Antibody mix 2 comprised different monoclonal antibodies to distinguish absolute numbers of γδ and non-γδ T cell subpopulations and to quantify CD25 activation marker expression on these cells in infected and mock-infected animals. The antibody combination in mix 3 was used to identify absolute counts of αβ T cell subsets and to phenotypically characterize the non-γδ T cells in more detail. Antibody mix 4 was used to examine the percentage of NK-like cells in PBMC, spleen and cecum of infected wild-type and γδ T cell knockout chickens only.

**Table 1.**
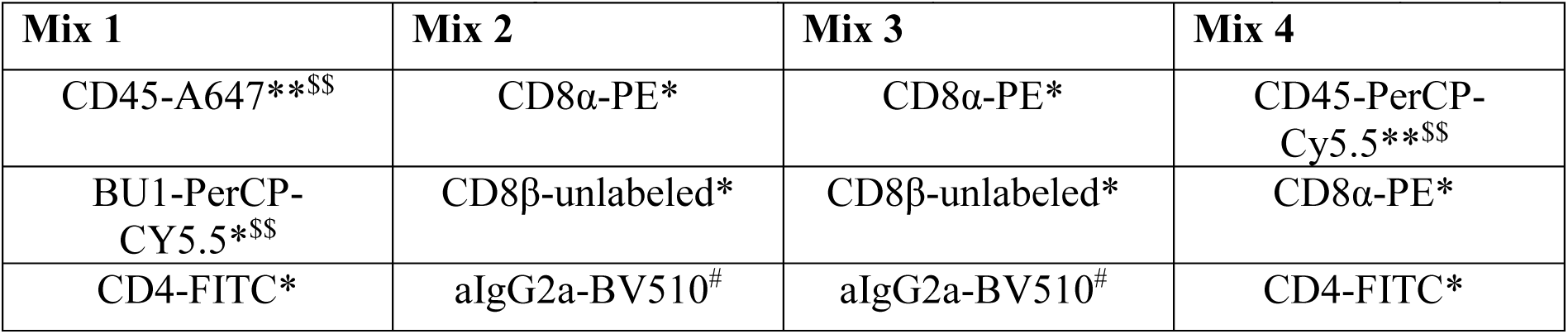
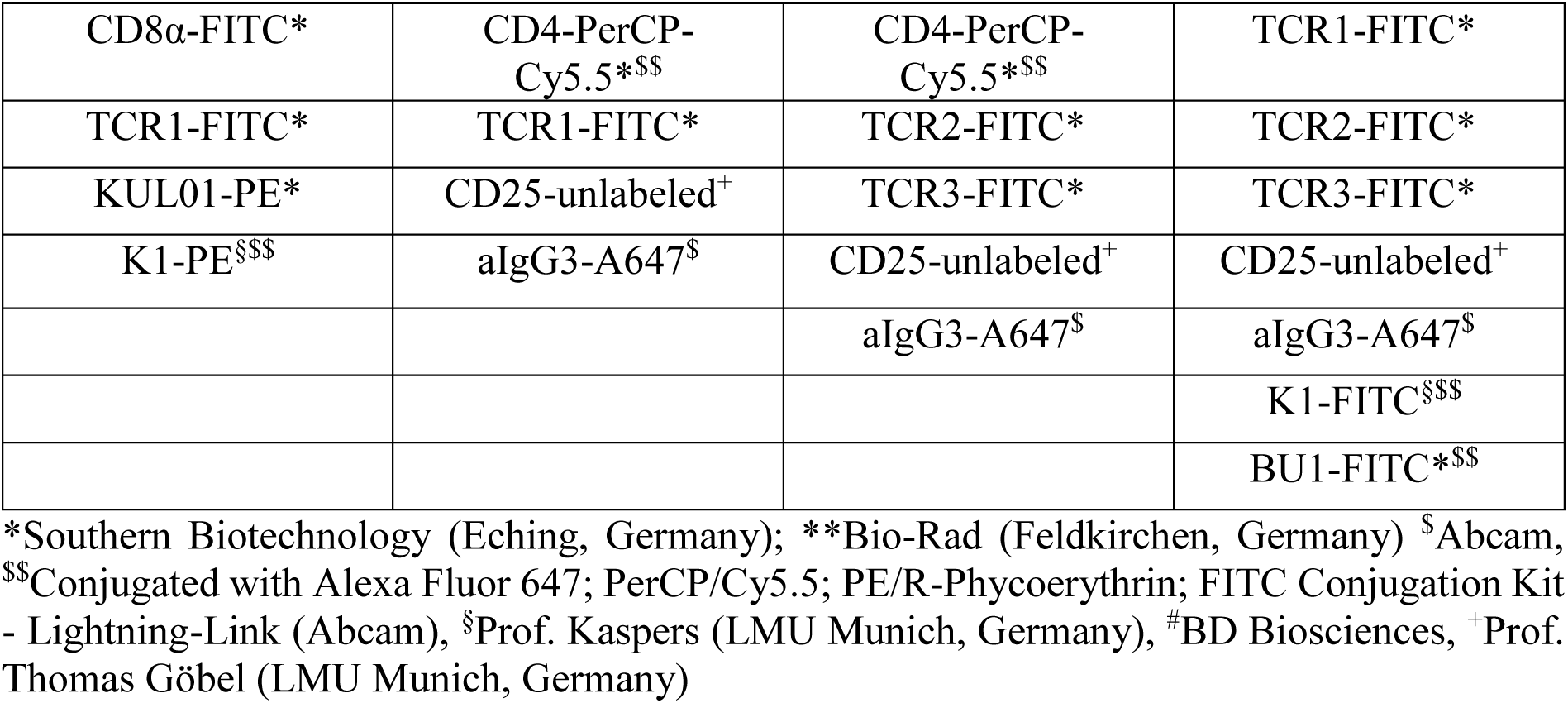
Overview of antigens labelled by antibody mixtures for flowcytometry analysis.

For the flow cytometric determination of the absolute immune cell numbers in whole blood, spleen and cecum, 20 µL of the whole blood samples were diluted in 980 µL PBS^+^ and used for cell labelling. Fifty microliters of the diluted whole blood or 50 µl of isolated immune cell suspensions from spleen and cecum were mixed with 20 µL of the appropriate monoclonal antibody combination (antibody mixes 1-3, Table 1) directly into Trucount tubes (BD Biosciences).

To assess the relative number of NK-like cells, 50 µL of the isolated immune cells from PBMC, spleen and cecum were incubated with 20 µL of the antibody mix 4 in a FACS tube without beads (Falcon^®^, Corning Inc., Corning, NY, USA).

All samples were incubated in the dark for 60 min. Shortly before flow cytometric analysis, 300 µL PBS^+^ and DAPI (1 µg/mL; Sigma, Taufkirchen, Germany) were added to each sample. Measurements were performed on a FACSCanto II (BD Biosciences) equipped with 488 nm, 633 nm and 405 nm lasers. Prior to the study, optimal antibody concentrations and compensations were determined for each antibody in the different mixes. Appropriate fluorescence-minus one controls were conducted to confirm specificity of antibody binding. Data acquisition was performed using the FACSDiva v9.0.1 software (BD Biosciences). For absolute cell analysis either 10,000 (mix 1) or 30,000 (mix 2 and 3) Trucount beads were acquired together with the cells. For mix 1, all CD45^+^ (UM16-6, IgG2a) cells as well as thrombocytes (K1^+^, IgG1; Prof. Kaspers) and monocytes (KUL01^+^, IgG1) were measured and data stored (58). For the mixes 2 and 3, the storage gate contained all γδ T cells (TCR1^+^, IgG1) and non-γδ T cells (TCR1^-^; mix 2), Vβ1^+^ and Vβ2^+^ T cells (TCR2^+^ and TCR3^+^, IgG1; mix 3) as well as the low and high-positive CD8α^+^ (CT-8, IgG1), CD8β^+^ (EP42, IgG2a), CD4^+^ (CT-4, IgG1) and CD25^+^ (28-4, IgG3) T lymphocytes. For mix 4, 30,000-150,000 (PBMC), 30,000-50,000 (spleen) and 10,000-30,000 (cecum) CD45^+^ cells were acquired and all events recorded per sample.

Data analysis was performed by using the FlowJo software (version 10, FlowJo, BD Biosciences). Before immune cell analysis, doublets were excluded (FSC-A versus FSC-W) and viable cells selected (DAPI-negative cells).

For mix 1, absolute numbers of thrombocytes and monocytes were measured from the K1/KUL01 dot plot against CD45. CD45-positive leukocytes were further used to distinguish Bu1-positive B lymphocytes (AV-20) and TCR1/CD4/CD8-positive T lymphocytes. Blood heterophils were selected by back-gating from the CD45^+^ versus SSC dot plot (58, 60).

In mix 2, distinct subsets of T cells were identified based on their expression or absence of the TCR1 marker (61). These included the CD8α^-^TCR1^+^, CD8α^++^TCR1^+^ and TCR1^-^ cell populations, as depicted on the CD8α versus TCR1 dot plot. The TCR1^-^ cells were further separated into CD8α^++^CD4^-^ and CD8α^+^CD4^+^ T cell subsets by plotting CD8α against CD4. The CD8α^++^TCR1^+^ and CD8α^++^CD4^-^TCR1^-^ T cell populations were further differentiated into the CD8αα homo- and CD8αβ heterodimer subsets. All lymphocyte populations were back-gated against SSC/FSC (Fig. S1).

For mix 3, T cell populations were gated based on their TCR2 and TCR3 expression as described for mix 2, respectively. For the CD8αα^++^CD4^-^TCR1^-^, CD8αα^++^TCR1^+^ and CD8αα^++^CD4^-^TCR2/3^+^ T lymphocyte subpopulations of the mixes 2-3, the absolute numbers of CD25-expressing cells were additionally determined.

Absolute cell numbers were calculated using following equation:

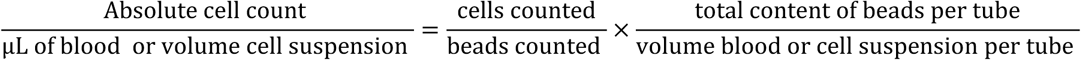

Absolute cell counts of isolated tissue cells were calculated for 1 mL and at the appropriate dilution and expressed as cells per 100 mg of spleen or per 10 cm^2^ of cecum.

In mix 4, the relative frequency of CD8α^+^ leukocytes (CD45^+^) lacking lineage-specific markers for T or B cells, referred to in this study as NK-like cells, was determined by plotting CD8α against CD45 (62). The lymphocyte subpopulation was confirmed by back-gating using an SSC/FSC dot plot. Cells expressing lineage-specific markers for TCR^+^ cells, B cells, CD4^+^ T cells, monocytes and thrombocytes (detected with FITC-conjugated antibodies) were excluded from the CD8α^+^CD45^+^ subset, as illustrated on CD8α versus FITC dot plots. Within the CD8α^+^CD45^+^FITC^-^ subpopulation, the relative proportion of CD25-expressing cells was additionally quantified to evaluate activation status of NK-like cells (Fig. S2).

### 2.6 Fluorescence-Activated Cell Sorting of T cell subsets

To investigate the transcriptional activity of various immune-related genes in T cells, fluorescence-activated cell sorting (BD FACS Aria II, BD Biosciences) was used with a 70 μm nozzle. We collected γδ (TCR1^+^) and non-γδ T cell (TCR1^-^) subsets from PBMC of *Salmonella*-infected chickens at 7, 9 and 12 dpi. Isolated blood cells (PBMC) were incubated with the specific monoclonal antibodies (CD8α-PE, CD8β-unlabeled, anti-IgG2a-BV510, CD4-PerCP-Cy5.5, TCR1-FITC) for 30 min in the dark and immediately sorted. Prior to sorting, the drop delay was adjusted using FACS AccuDrops (BD Biosciences). A test sort was performed to ensure a purity level exceeding 95% in the sorted cell subsets. CD8αα^++^ and CD8αβ^++^ γδ T cells as well as CD8αα^++^CD4^-^ and CD8αβ^++^CD4^-^ non-γδ T cell subsets were collected separately. The sorted cell populations from each bird were immediately stored in RNAlater (Qiagen) at −20 °C until RNA isolation.

### 2.7 Immunohistochemistry

To compare the *Salmonella* invasion of the wild-type and TCR Cγ^−/−^ chickens, frozen sections of the cecum of *Salmonella*-infected chickens were prepared. The immunohistochemical staining method has been described previously (9). Briefly, tissue sections (7 µm thickness) were fixed in acetone and subsequently incubated with the monoclonal antibodies against *Salmonella enterica* lipopolysaccharide (clone 5D12A 1:800 in PBS, Bio-Rad, Feldkirchen, Germany). Next, the peroxidase-anti-peroxidase conjugated secondary goat-anti-mouse immunoglobulin (Sigma, Deisenhoven, Germany) was applied. The enzyme-linked antibody was visualized by reaction with 3,3’-diaminobinzidine (DAB, Merck, Darmstadt, Germany) and hydrogen peroxide. The sections were counterstained with haematoxylin (Agilent Dako, Waldbronn, Germany) and finally mounted with Canada balsam (Carl Roth, Karlsruhe, Germany). As a negative control, the primary antibodies were replaced by PBS.

### 2.8 Image analysis

The *Salmonella* invasion into the cecal mucosa was determined by analyzing the percentage of *Salmonella* positively-stained areas in the cecal epithelial lining and lamina propria of infected γδ T cell knockout and wild-type chickens. Analysis was performed using the image analysis system cellSens Dimension software (Olympus, Hamburg, Germany) as previously described (9). Regions of interest (ROIs) were drawn and at least 1 mm^2^ of epithelium and 1 mm^2^ of lamina propria analyzed per animal.

### 2.9 Quantitative RT-PCR for chemokine, cytokine and iNOS expression

To investigate the transcriptional dynamics of important immune genes after *Salmonella* infection, total RNA was extracted from cecal tissue and sorted cells using the RNeasy Mini Kit (Qiagen, Hilden, Germany) according to the manufacturer’s protocol. The RNase-free DNase kit (Qiagen) was used to digest any residual DNA. Quantitative RT-PCR was carried out using QuantiFast^®^ SYBR^®^ Green PCR kit (Qiagen), as described by the manufacturer and in previous studies (9, 20). Sequences, annealing temperatures and accession numbers of the primers for IFN-γ, inducible nitric oxide synthase (iNOS; (63); cecum only), glycerinaldehyd-3-phosphat-dehydrogenase (GAPDH), lipopolysaccharide-induced tumor necrosis factor-alpha factor (LITAF; (9); cecum only), IL-10 (cecum only), IL-1β (cecum only), IL-17A, IL-22 (64), IL-2Rα (20) have been previously described. Newly designed primers used in this study are provided in Table 2. The expression of target genes was normalized to the housekeeping gene glycerinaldehyde-3-phosphate (GAPDH). Results are presented as 40-ΔCt values.

**Table 2.**
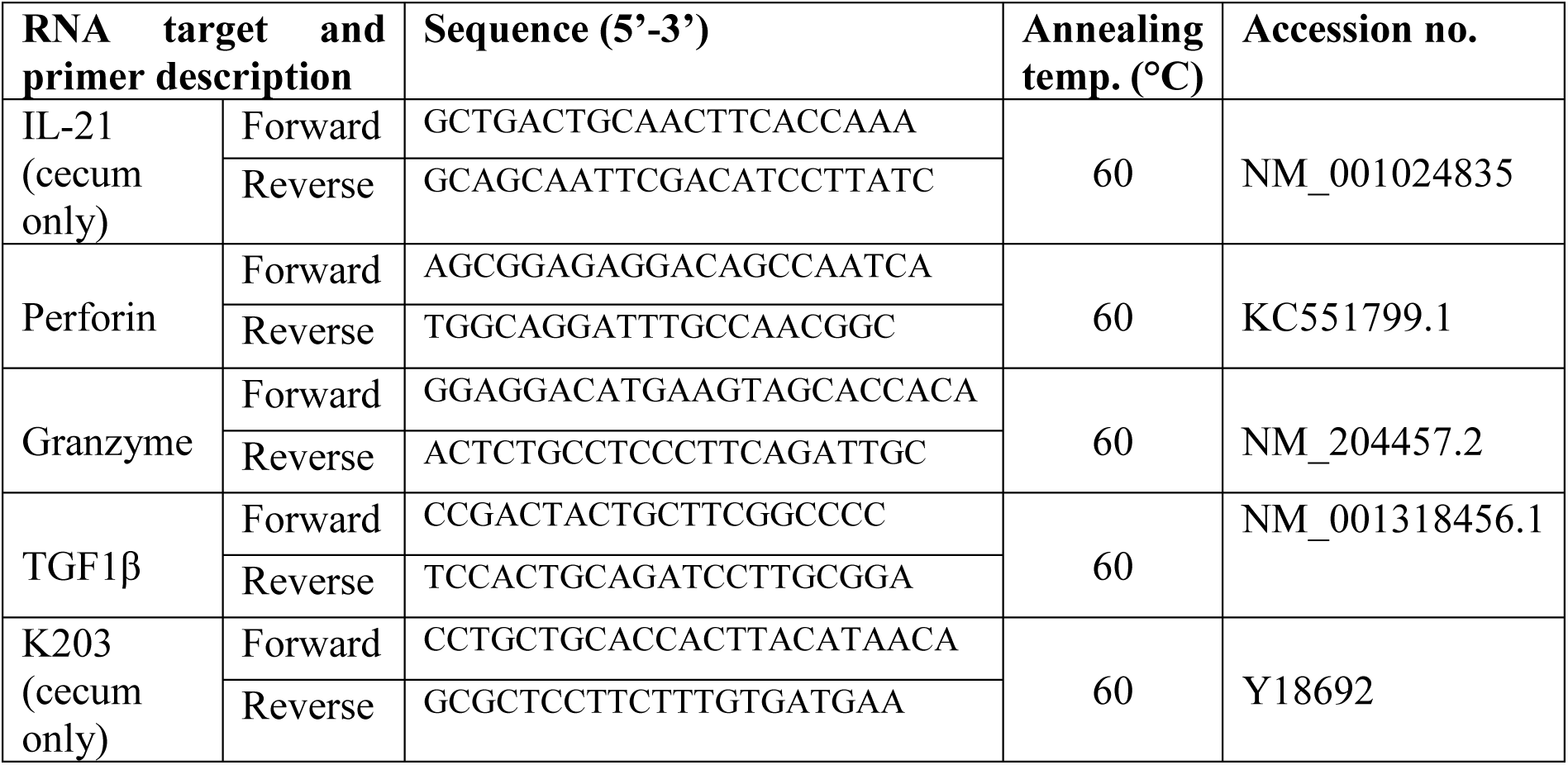
Primer pairs used for gene expression analysis.

### 2.10 Statistical analysis

For analysis of microbiological data, viable bacterial counts were converted into logarithmic form. Samples with a viable count below the detection limit for direct plating (log_10_ < 1.47) but positive after enrichment were assigned a value of log_10_ = 1.0. Samples with no detectable *Salmonella* growth after enrichment was assigned a log_10_ value of 0. Data were evaluated using multifactorial variance analysis with group and time as factors. P values < 0.05 were considered as statistically significant (software: Statgraphics Plus, Inc., Rockville, MD, USA).

Normally distributed data of flow cytometry, qPCR and histochemistry (Shapiro-Wilk test < 0.05) were analyzed by 2-way ANOVA followed by Tukey’s multiple comparison test (GraphPad Prism 7, GraphPad Software, Inc, Boston, MA, USA). Mann-Whitney U test was applied for not normally distributed data. Infected and mock-infected chickens as well as wild-type and γδ T cell knockout animals were compared to each other at multiple time points. P values < 0.05 were considered significant.

## 3. Results

### 3.1 Increased numbers of Salmonella in liver of TCR Cγ^−/−^ chickens compared to wild-type chickens

To examine the propagation of *S*. Enteritidis in infected wild-type and TCR Cγ^−/−^ chickens, the cfu of the bacteria were determined in liver and cecal content.

The oral administration of 2 x 10^7^ cfu of *S*. Enteritidis per bird resulted in robust cecal colonization and systemic spread into the liver, without inducing clinical signs. Notably, at one day post infection (dpi), we observed equivalent high *Salmonella* loads in the cecal content of both chicken lines, indicating a comparable infectious pressure at the beginning of the disease (Fig. 1A). No significant differences in cfu were detected in cecal contents of wild-type and TCR Cγ^−/−^ chickens at later stages of infection.

**Figure 1:**
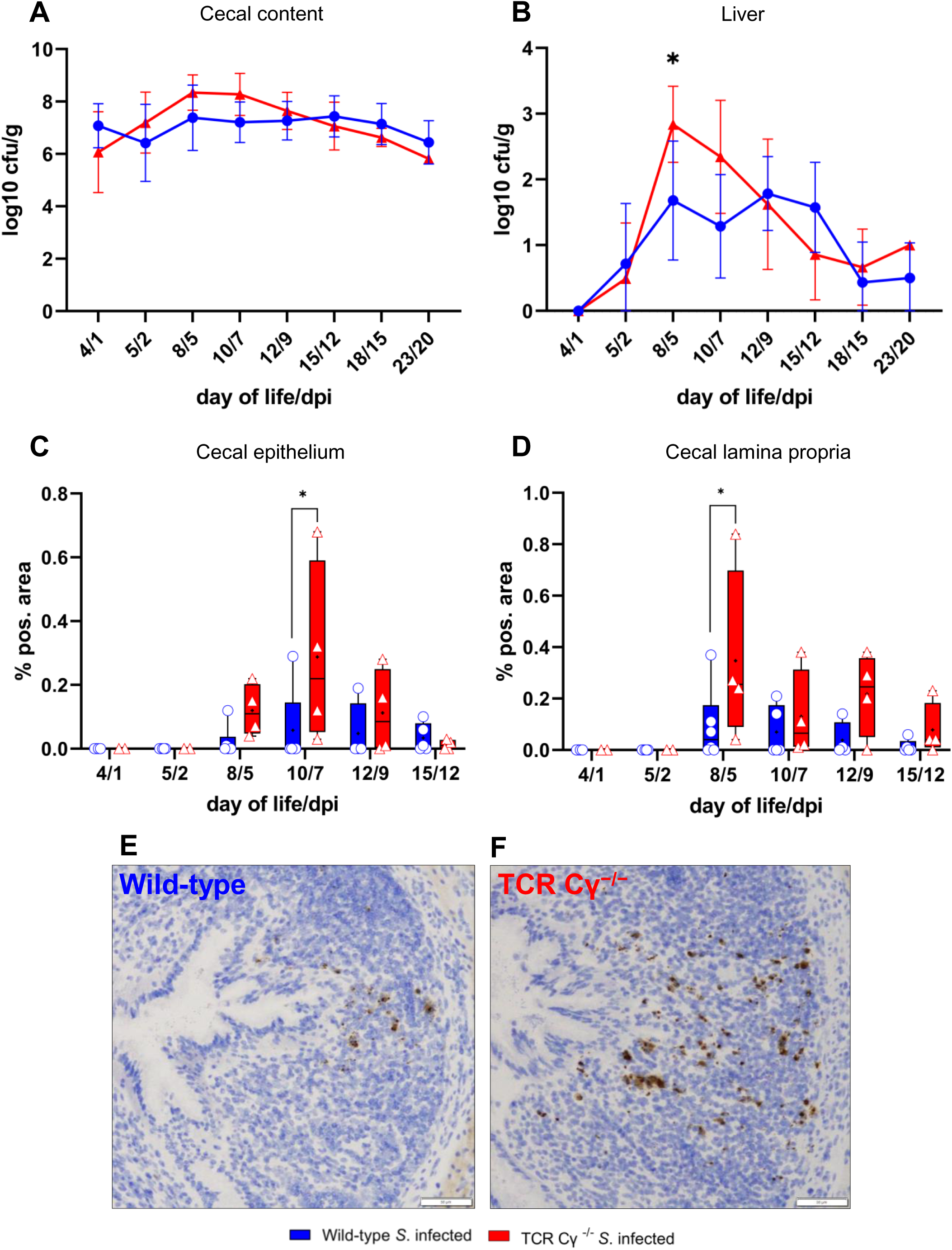
Detection of *Salmonella* Enteritidis in wild-type and TCR Cγ^−/−^ chickens post infection. Logarithmic bacterial counts of *Salmonella* in cecal content (A) and liver (B) of wild-type and TCR Cγ^−/−^ chickens. Data represent the mean ± standard deviation; n = 3-9. Box plot diagrams displaying the percentage of *Salmonella*-positive area in cecal epithelium (C) and in lamina propria (D) of infected wild-type and TCR Cγ^−/−^ chickens. Data are presented as minimum and maximum cell counts, with median indicated; n = 2-6. Representative immunohistochemical staining of *Salmonella* (brown) in cecal tissue from infected wild-type (E) and TCR Cγ^−/−^ chickens (F) at 5 dpi. Scale bar indicates 50 µm. * indicates significant differences between chicken groups, p < 0.05.

In the liver, significant differences in *S*. Enteritidis colonization were observed between the genetically different chicken lines. Compared to wild-type birds, significantly higher cfu of *Salmonella* were found in the liver of TCR Cγ^−/−^ chickens at 5 dpi (Fig. 1B). The dynamics of *Salmonella* counts in the liver also differed between the two chicken lines during the course of the infection. In the absence of γδ T cells, the number of cfu in the liver increased rapidly after infection, peaking at 5 dpi. Thereafter, the bacterial load decreased quickly. In contrast, the *Salmonella* load in the liver of wild-type animals remained relatively low and stable throughout the infection. At 15 and 20 dpi, hardly any *Salmonella* were detectable in livers of wild-type and knockout chickens. No *Salmonella* was detected in the liver or cecum of mock-infected chickens at any time point.

### 3.2 Increased Salmonella invasion into the cecum wall of TCR Cγ^−/−^ chickens compared to wild-type chickens

The invasion of *S*. Enteritidis into the cecal wall was assessed using immunohistochemistry and image analysis. The results demonstrated that *Salmonella* was able to invade the cecal mucosa in both chicken lines, albeit at different levels. At 7 dpi, a significantly higher percentage of *Salmonella*-positive areas were observed in the cecal epithelium of TCR Cγ^−/−^ chickens compared to wild-type birds. A trend towards increased *Salmonella* invasion into the intestinal epithelium was also noted in the absence of γδ T cells at 5 and 9 dpi (Fig. 1C).

In the lamina propria of γδ T cell knockout chickens, a significantly increased percentage of *Salmonella* was detected at 5 dpi, with a trend towards an increased percentage of *Salmonella*-positive areas also observed at 9 dpi in comparison to infected wild-type chickens (Fig. 1D-F).

### 3.3 Elevated numbers of monocytes in blood of wild-type and TCR Cγ^−/−^ chickens after Salmonella infection

Dynamic changes in absolute leukocyte numbers in whole blood of *Salmonella*-infected γδ T cell knockout and wild-type chickens were quantified by flow cytometry (Fig. 2). Compared with mock-infected controls, *S.* Enteritidis exposure induced a rapid and significant increase in monocytes at 5 dpi in both wild-type and TCR Cγ^−/−^ chickens, with elevated levels persisting until at least 12 dpi in both chicken lines. In the absence of γδ T cells, knockout chickens displayed a trend towards higher monocyte counts at 5 and 7 dpi compared to wild-type birds, accompanied by large individual differences. Additionally, a significant increase in circulating thrombocyte numbers in both infected wild-type and TCR Cγ^−/−^ chickens was observed compared to mock controls. No relevant differences were detected in the counts of heterophils and B cells, whereas TCR Cγ^−/−^ chickens showed a trend towards lower blood T cell numbers (Fig. S3).

**Figure 2:**
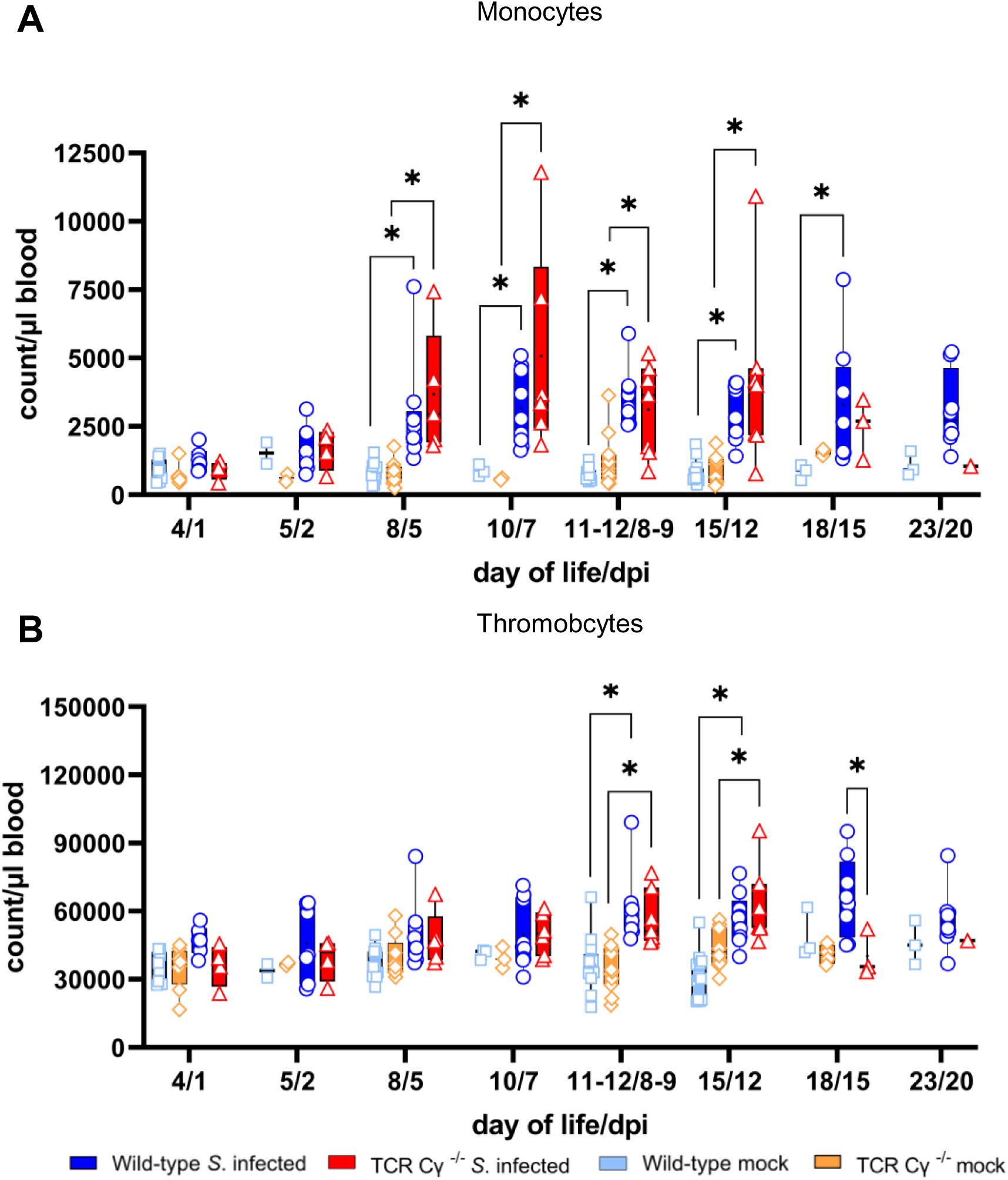
Flow cytometric analysis of leukocytes in avian whole blood following *Salmonella* Enteritidis infection. Absolute numbers of viable monocytes (A) and thrombocytes (B) were measured over time in blood from wild-type and TCR Cγ^−/−^ chickens following *Salmonella* or mock infection. Data are presented as minimum and maximum cell counts, with median indicated; n = 1-13. * indicates significant differences between chicken groups, p < 0.05. No data available for mock-infected TCR Cγ^−/−^ chickens at 20 dpi.

### 3.4 Absolute numbers of γδ T cell subsets increased in whole blood, spleen and cecum after Salmonella *infection of wild-type chickens*

Determination of absolute numbers of γδ T cell subsets (TCR1^+^) in whole blood, spleen and cecum revealed an increase of CD8αα^++^ γδ T cells across all compartments (Fig. 3A). In whole blood, significantly enhanced numbers of CD8αα^++^ γδ T cells were observed at 5 dpi and between 8-9 and 15 dpi. At 8-9 and 12 dpi, the number of CD8αα^++^ γδ T cells co-expressing the CD25 antigen also increased significantly (Fig. 3B). In the spleen of *Salmonella*-infected wild-type chickens, significantly elevated numbers of CD8αα^++^ γδ T cells were detected at 9 and 15 dpi. A significant increase in CD8αα^++^CD25^+^ γδ T cells was observed at 9 dpi, with a similar trend at 12 and 15 dpi. Comparable results were found in the cecum, where CD8αα^++^ γδ T cells increased significantly at 9 dpi, with a similar trend noted at 7, 12 and 15 dpi compared to mock-infected controls.

**Figure 3:**
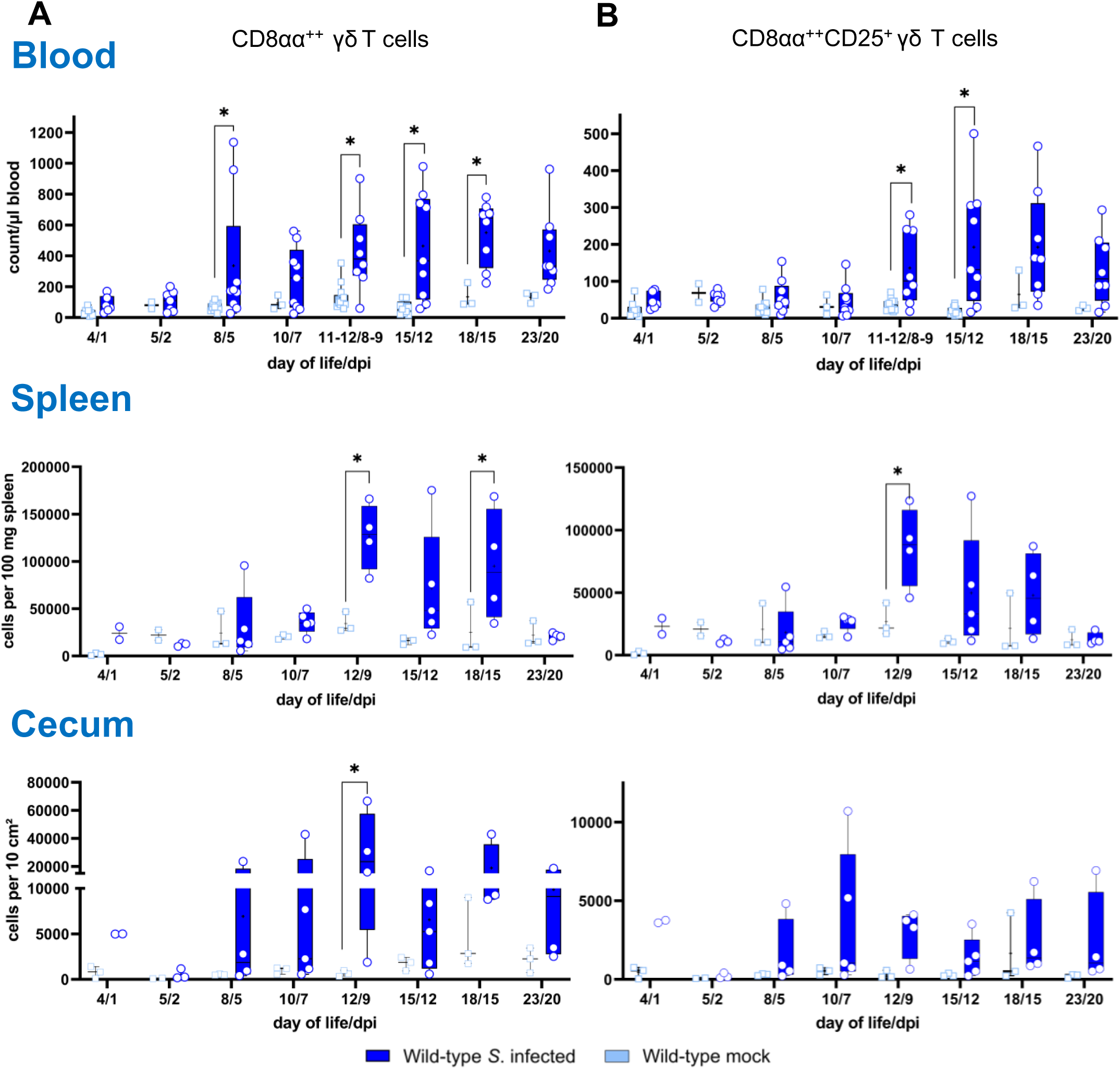
Flow cytometric analysis of γδ T cell subsets in wild-type chickens after *Salmonella* Enteritidis infection. Absolute numbers of CD8αα^++^ (A) and CD8αα^++^CD25^+^ (B) γδ T cells are shown of blood (n = 2-13), spleen (n = 2-5) and cecum (n = 2-5) from *Salmonella*-infected and mock-infected wild-type chickens. Data are presented as minimum and maximum cell counts per μL blood, per 100 mg spleen and per 10 cm² cecum, with medians indicated. * indicates significant differences between chicken groups, p < 0.05.

### 3.5 Compensation of the lack of γδ T cells in TCR Cγ^−/−^ chickens after Salmonella infection

Flow cytometric analysis revealed an increased emergence of a CD8αα^++^CD4^-^ non-γδ T cell population in TCR Cγ^−/−^ chickens (Fig. 4). Following *Salmonella* infection, the absolute numbers of these cells were significantly increased in peripheral blood and spleen in the absence of γδ T cells (8-9 and 12 dpi), but not in wild-type birds (Fig. 4A). Moreover, higher absolute numbers of a CD8αα^++^CD4^-^CD25^+^ non-γδ T cell phenotype were found in blood (8-9 and 12 dpi) and spleen (9 dpi) of TCR Cγ^−/−^ chickens compared to wild-type (Fig. 4B). Notably, these non-γδ T cells identified in the infected TCR Cγ^−/−^ chickens exhibited an identical sub-phenotype (CD8αα^++^ and CD25^+^) as the γδ T cells that were elevated in infected wild-type chickens (Fig. 3). Analysis of cecal tissue showed no significant differences between the chicken groups.

**Figure 4:**
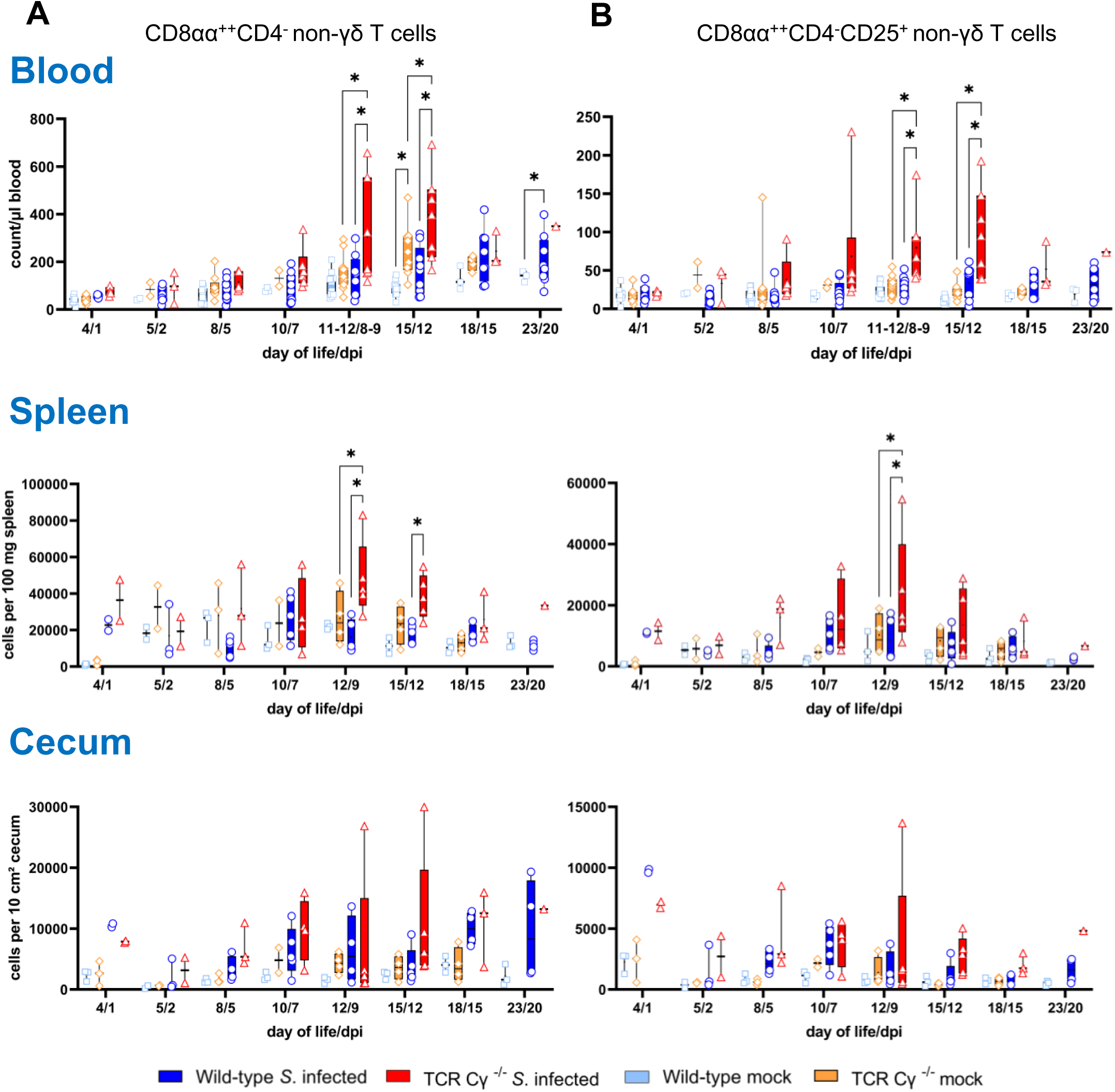
Flow cytometric analysis of non-γδ T cell subsets in wild-type and TCR Cγ^−/−^ chickens after *Salmonella* Enteritidis infection. Absolute numbers of CD8αα^++^CD4^-^ (A) and CD8αα^++^CD4^-^CD25^+^ (B) non-γδ T cells are shown for blood (n = 1-13), spleen (n = 1-5) and cecum (n = 1-5) from *Salmonella*-infected and mock-infected wild-type and TCR Cγ^−/−^ chickens. Data are presented as minimum and maximum cell counts per μL blood, per 100 mg spleen and per 10 cm² cecum, with medians indicated. * indicates significant differences between chicken groups, p < 0.05. No data available for mock-infected TCR Cγ^−/−^ chickens at 20 dpi.

### 3.6 CD8αα^++^CD4^-^ non-γδ T cells of Salmonella-infected TCR Cγ^−/−^ chickens consist at least partially of αβ T cells

To further characterize the phenotype of the CD8αα^++^CD4^-^ non-γδ T cells observed in TCR Cγ^−/−^ animals, the emergence of Vβ1 (TCR2)/Vβ2 (TCR3) αβ T cells in blood and spleen was investigated, relative to the number of non-γδ T cells in these compartments (Fig. 5). In the spleen, a significant increase in Vβ1/Vβ2 cells expressing CD8αα^++^ was detected in TCR Cγ^−/−^ chicken at 9 dpi compared to both infected wild-type and mock-infected TCR Cγ^−/−^ controls, with a similar trend observed at 12 and 15 dpi (Fig. 5A). Notably, this elevated population of Vβ1/Vβ2 αβ T cells did not fully account for the entire increase of CD8αα^++^CD4^-^ non-γδ T cells observed in spleens of TCR Cγ^−/−^ chickens (Fig. 5B).

**Figure 5:**
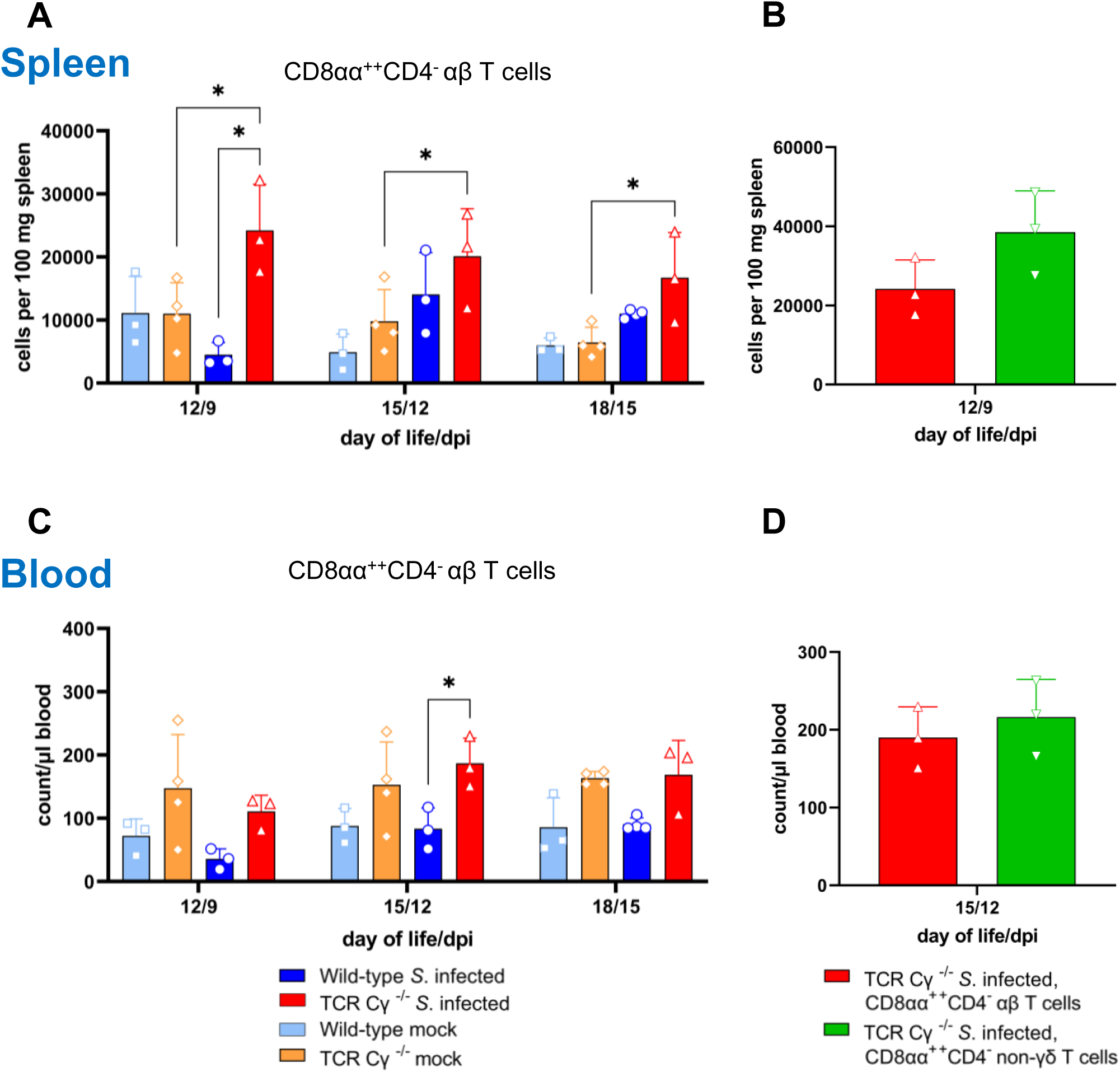
Emergence of CD8αα^++^CD4^-^ αβ T cells in TCR Cγ^−/−^ chickens following *Salmonella* Enteritidis infection. Absolute numbers of CD8αα^++^CD4^-^ Vβ1/Vβ2 αβ T cell subsets are shown for spleen (A) and blood (C) of *Salmonella*-infected and mock-infected wild-type and TCR Cγ^−/−^ chickens. The number of CD8αα^++^CD4^-^ Vβ1/Vβ2 αβ T cells are compared to the CD8αα^++^CD4^-^ non-γδ T cell population in spleen (B) and blood (D) of the same *Salmonella*-infected TCR Cγ^−/−^ chickens. Data represent the mean ± standard deviation; n = 3-4. * indicates significant differences between chicken groups, p < 0.05.

In blood, the findings were consistent with those in the spleen. A significant increase in CD8αα^++^CD4^-^ αβ T cells was detected in infected TCR Cγ^−/−^ birds at 12 dpi compared to infected wild-type animals. However, no significant differences were observed compared to mock-infected TCR Cγ^−/−^ controls (Fig. 5C-D).

### 3.7 Increase of CD8α^+^CD4^+^ non-γδ T cells in TCR Cγ^−/−^ chickens and wild-type chickens after Salmonella infection

Analysis of CD4^+^ T cell populations in experimental chickens revealed a significant expansion in the absolute numbers of CD8α^+^CD4^+^ (Fig. S4A) and CD8α^+^CD4^+^CD25^+^ T cells (Fig. S4B) in avian peripheral blood after *Salmonella* infection. This increase was prominent between 8-9 and 12 dpi in γδ T cell knockout and wild-type chickens. Notably, significant elevated numbers of CD8α^+^CD4^+^ (15 dpi) or CD8α^+^CD4^+^CD25^+^ T cells (7, 9, 12 and 15 dpi) were detected in the spleen of infected chickens. In cecum, the number of CD8α^+^CD4^+^ T cells was increased only in wild-type chickens at 15 dpi.

### 3.8 Higher percentages of NK-like cells expressing CD25 in blood of TCR Cγ^−/−^ than wild-type chickens after Salmonella infection

To further identify potential compensatory mechanisms in the absence of γδ T cells, the occurrence of NK-like cells in PBMCs, spleen, and cecum of chicken following *Salmonella* infection was analyzed. Our findings showed a low relative abundance of NK-like cells in PBMCs and spleen (2–8%), whereas a notably higher proportion (30–40%) of NK-like cells was detected in the cecum of both wild-type and TCR Cγ^−/−^ chickens (Fig. 6A). No significant differences in the frequency of NK-like cell were observed between the two *Salmonella*-infected animal groups. However, the activation status of the cells showed a significant increase of CD25^+^ NK-like cells in PBMCs of TCR Cγ^−/−^ chickens compared to wild-type birds between 9 and 12 dpi (Fig. 6B). In chickens of both genotypes, a high proportion of NK-like cells in the cecum demonstrated a CD25 expression (activated phenotype) which decreased over time. Nevertheless, a trend towards more CD25^+^ NK-like cells was obvious in the spleen and cecum of TCR Cγ^-/-^ compared to wild-type chickens between 9 and 15 dpi.

**Figure 6:**
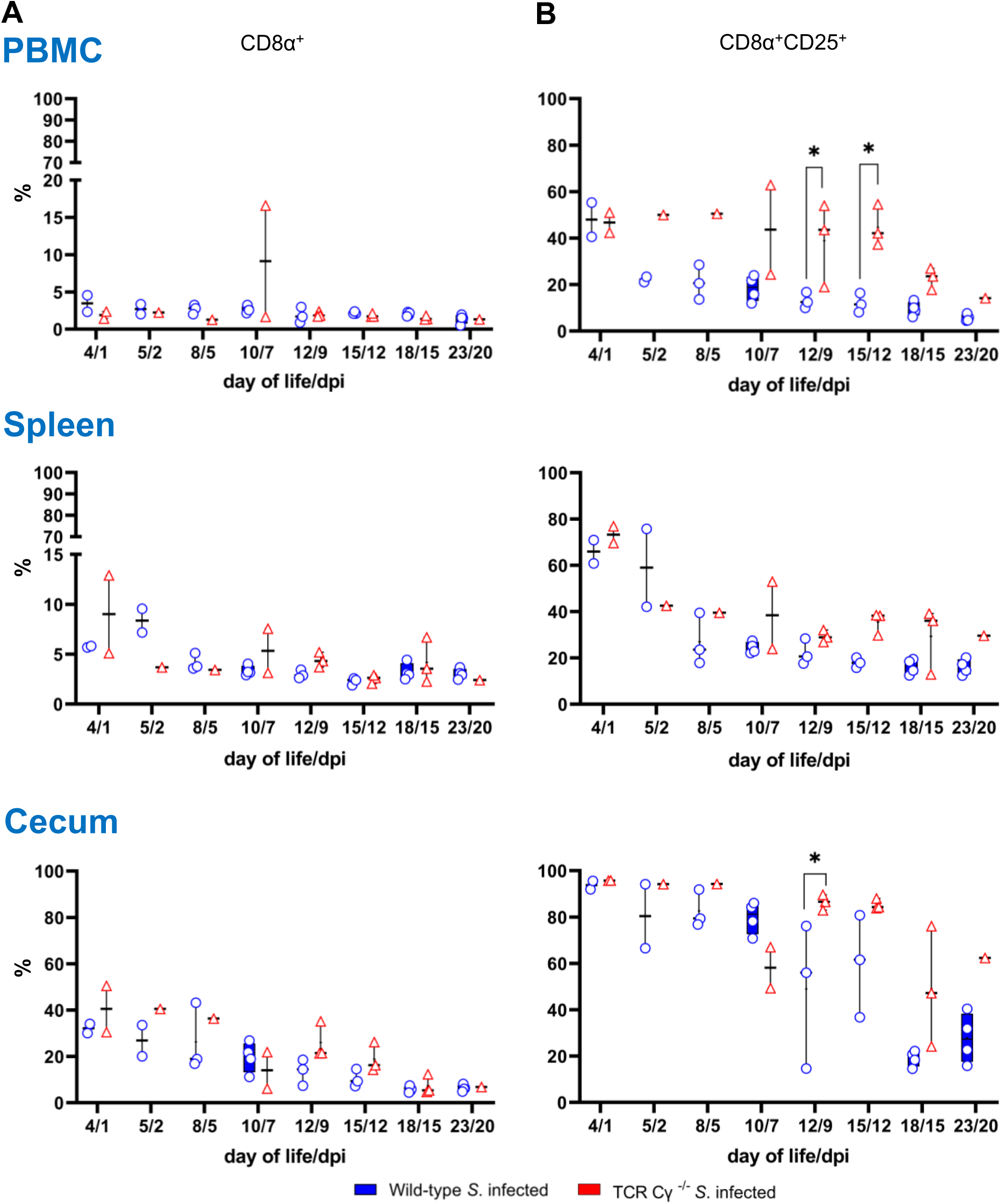
Frequency and activation status (CD25 expesssion) of NK-like cells in wild-type and TCR Cγ^−/−^ chickens after *Salmonella* Enteritidis infection. (A) Relative frequency of CD8α^+^ NK-like cells within the CD45^+^ lymphocyte population lacking T and B cell lineage as well as monocyte markers (TCR1, TCR2, TCR3, CD4, Bu1, K1) in PBMCs, spleen and cecum of *Salmonella*-infected wild-type and TCR Cγ^−/−^ chickens. (B) Percentage of CD8α^+^ NK-like cells expressing CD25 (marker of cell activation) relative to all CD8α^+^ NK-like cells. Data are presented as minimum and maximum cell counts, with median indicated; n = 1-4. * indicates significant differences between chicken groups, p < 0.05.

### 3.9 CD8αα^++^CD4^-^ non-γδ T cells of Salmonella-infected TCR Cγ^−/−^ chickens exhibit transcription levels indicative of a pre-activated state

To compare functional characteristics of CD8αα^++^CD4^-^ non-γδ T cells from TCR Cγ^−/−^ birds with the CD8αα^++^CD4^-^ non-γδ T cell and CD8αα^++^ γδ T cell counterparts from wild-type chickens after infection, the transcription levels of immune-related genes in sorted cells were analyzed by quantitative real-time RT-PCR.

Our results revealed that CD8αα^++^CD4^-^ non-γδ T cells from both *Salmonella*-infected wild-type and TCR Cγ^−/−^ chickens exhibited resembling gene expression profiles. No significant differences were observed for the transcription levels of IL-17A, IL-2Rα, IL-22, IFN-γ, TGFβ, perforin and granzyme (Fig. 7). This indicates that these cells are not only phenotypically but also functionally comparable. However, it should be noted that the CD8αα^++^CD4^-^ non-γδ T cells were present in significantly higher numbers in infected TCR Cγ^−/−^ chickens than in wild-type animals.

**Figure 7:**
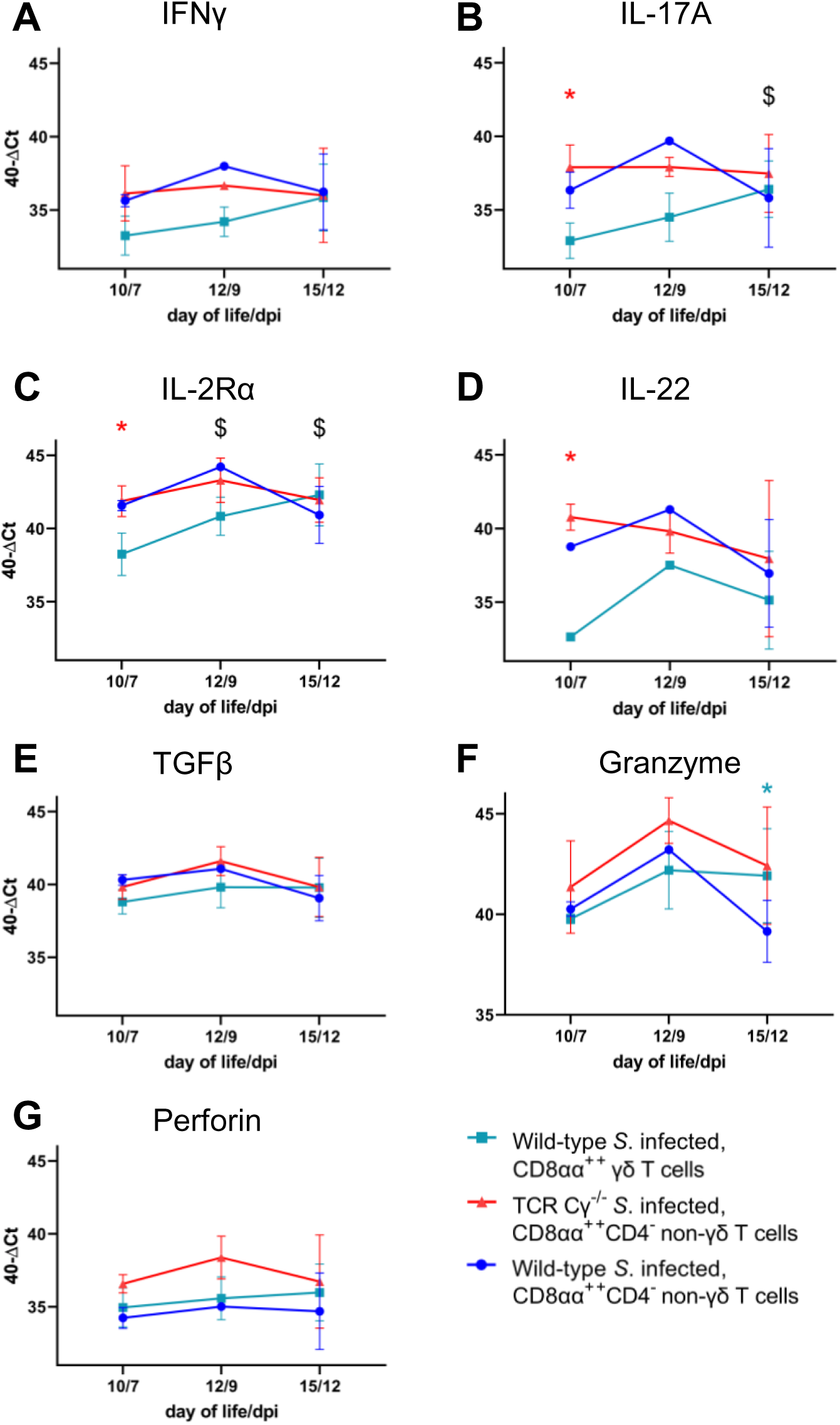
RT qPCR analysis of sorted CD8αα^++^ γδ and non-γδ T cells from *Salmonella* Enteritidis-infected wild-type and TCR Cγ^−/−^ chickens. Transcription levels of IFN-γ, IL-17A, IL-2Rα, IL-22, TGFβ, granzyme, and perforin (A-G) were measured in CD8αα^++^ γδ and CD8αα^++^CD4^-^ non-γδ T cells from wild-type chickens, as well as CD8αα^++^CD4^-^ non-γδ T cells from TCR Cγ^−/−^ animals. Data represent the mean 40-ΔCt ± standard deviation; n = 1-6. * indicates significant difference between CD8αα^++^ γδ T cells of wild-type chickens and CD8αα^++^CD4^-^ non-γδ T cells of TCR Cγ^−/−^ birds. * indicates a significant difference between CD8αα^++^ γδ T cells and CD8αα^++^CD4^-^ non-γδ T cells within wild-type chickens. $ indicates significant different transcription levels of CD8αα^++^ γδ T cells compared to 7 dpi, p < 0.05.

CD8αα⁺⁺CD4^-^ non-γδ T cells of infected TCR Cγ^−/−^ chickens exhibited significantly elevated transcription of IL-17A, IL-2Rα, and IL-22 at 7 dpi compared to CD8αα⁺ γδ T cells (always CD4^-^) of infected wild-type chickens. These transcriptional differences diminished by 12 dpi, driven by a significant increase in transcriptional activity in CD8αα⁺ γδ T cells over time (Fig. 7B-D). The same trend was observed for IFN-γ, but without significant differences (Fig. 7A). No significant transcriptional differences were detected for TGFβ or perforin between T cell subsets. However, granzyme expression was significantly increased in CD8αα^++^ γδ T cells at 12 dpi compared to their CD8αα^++^CD4^-^ non-γδ counterparts in infected wild-type birds (Fig. 7E-G).

### 3.10 Significant differences in gene transcription of IL-17A and IL-2Rα in CD8αα^++^CD4^-^ and CD8αβ^++^CD4^-^ non-γδ T cells of Salmonella-infected TCR Cγ^−/−^ chickens

In *Salmonella*-infected TCR Cγ^−/−^ chickens, CD8αα⁺⁺CD4^-^ non-γδ T cells showed higher transcription of IL-17A (at 7 and 12 dpi) and IL-2Rα (at 7, and 12 dpi) compared to CD8αβ⁺⁺CD4^-^ non-γδ T cells, indicating the activated state of CD8αα^++^CD4^-^ non-γδ T cells (Fig. 8B-C). Moreover, IL-2Rα transcription was significantly increased in CD8αβ^++^ γδ T cells from infected wild-type chickens compared to CD8αβ^++^CD4^-^ non-γδ T cells in infected TCR Cγ^−/−^ birds at 7 and 12 dpi, suggesting distinct functional capabilities between these cell populations. Additionally, IL-2Rα was more highly transcribed in CD8αα⁺⁺ γδ T cells than CD8αβ⁺⁺ γδ T cells of infected wild-type chickens at 12 dpi.

**Figure 8:**
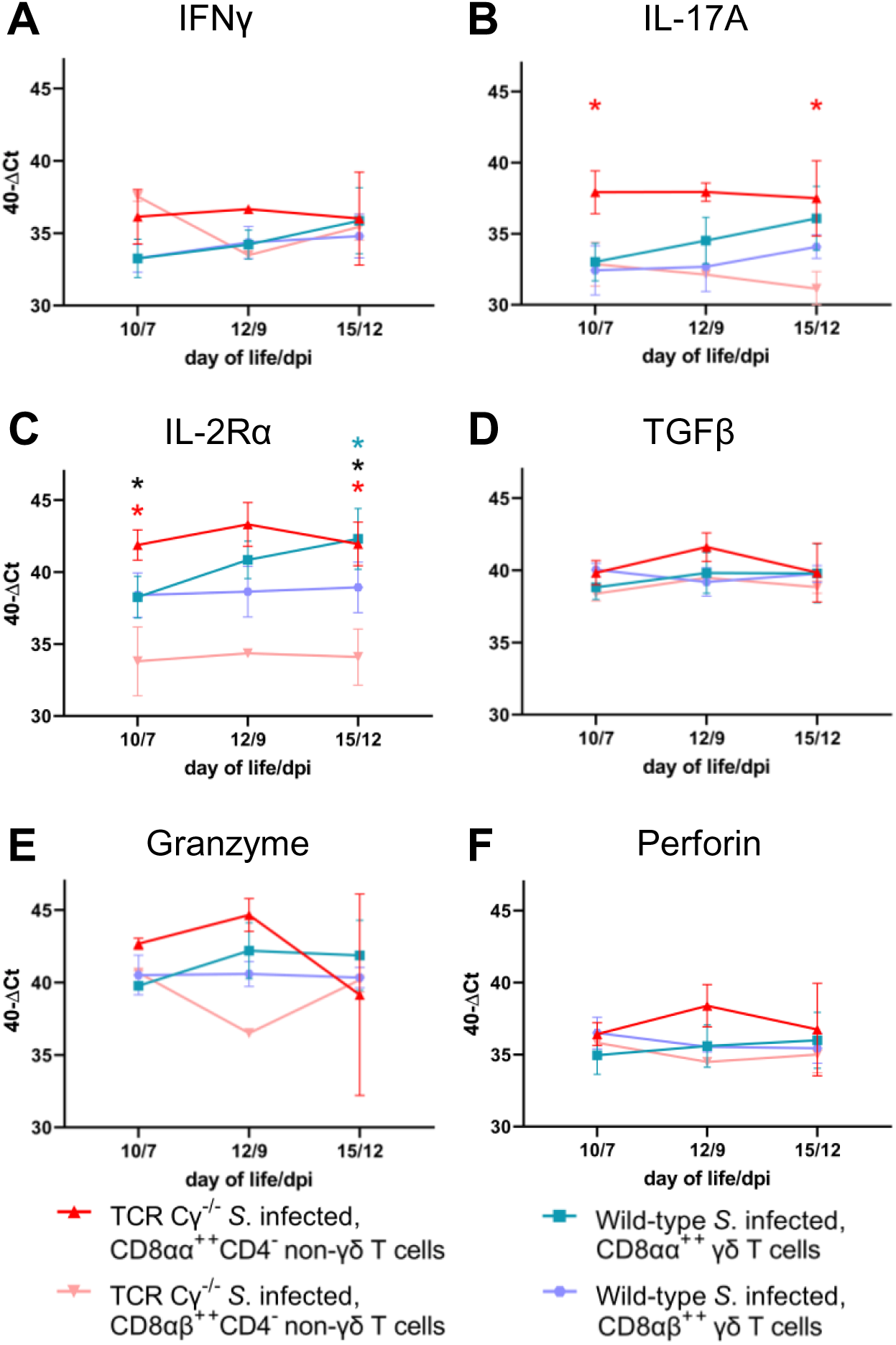
RT qPCR analysis of sorted CD8αα^++^ and CD8αβ^++^ T cell subsets from *Salmonella* Enteritidis-infected wild-type and TCR Cγ^−/−^ chickens. Transcription levels of IFN-γ, IL-17A, IL-2Rα, TGFβ, granzyme, and perforin (A-F) were measured in CD8αα^++^ and CD8αβ^++^ γδ T cells from wild-type chickens, as well as CD8αα^++^CD4^-^ and CD8αβ^++^CD4^-^ non-γδ T cells from TCR Cγ^−/−^ animals. Data represent the mean 40-ΔCt ± standard deviation; n = 1-6. * indicates significant difference between CD8αα^++^CD4^-^ and CD8αβ^++^CD4^-^ non-γδ T cells of TCR Cγ^−/−^ animals. * indicates significant difference between CD8αα^++^ and CD8αβ^++^ γδ T cells of wild-type animals. * indicates a significant difference between CD8αβ^++^CD4^-^ non-γδ T cells from TCR Cγ^−/−^ animals and CD8αβ^++^ γδ T cells from wild-type birds, p < 0.05.

### 3.11 Significant differences in gene transcription of IFN-γ and iNOS in cecal tissue of Salmonella-infected wild-type and TCR Cγ^−/−^ chickens

To elucidate the impact of γδ T cells on the temporal regulation of immune gene transcription during the course of infection, we assessed the transcriptional profiles of key cytokines in the cecum of the wild-type and TCR Cγ^−/−^ chickens.

Both wild-type and knockout chickens showed significant upregulation of pro-inflammatory cytokines after *Salmonella* infection, including IL-1β, IL-22, LITAF, and the chemokine K203, compared to mock-infected controls (Fig. 9). Upregulation generally occurred between 5 and 12 dpi. Notably, IFN-γ expression was significantly elevated at 2 dpi exclusively in TCR Cγ^−/−^ chickens, exceeding the levels observed in infected wild-type birds (Fig. 9A). Additionally, iNOS was upregulated in both groups at 5, 7 and 9 dpi, with TCR Cγ^−/−^ chickens demonstrating significantly higher iNOS transcription at 5 dpi compared to wild-type chickens (Fig. 9B).

**Figure 9:**
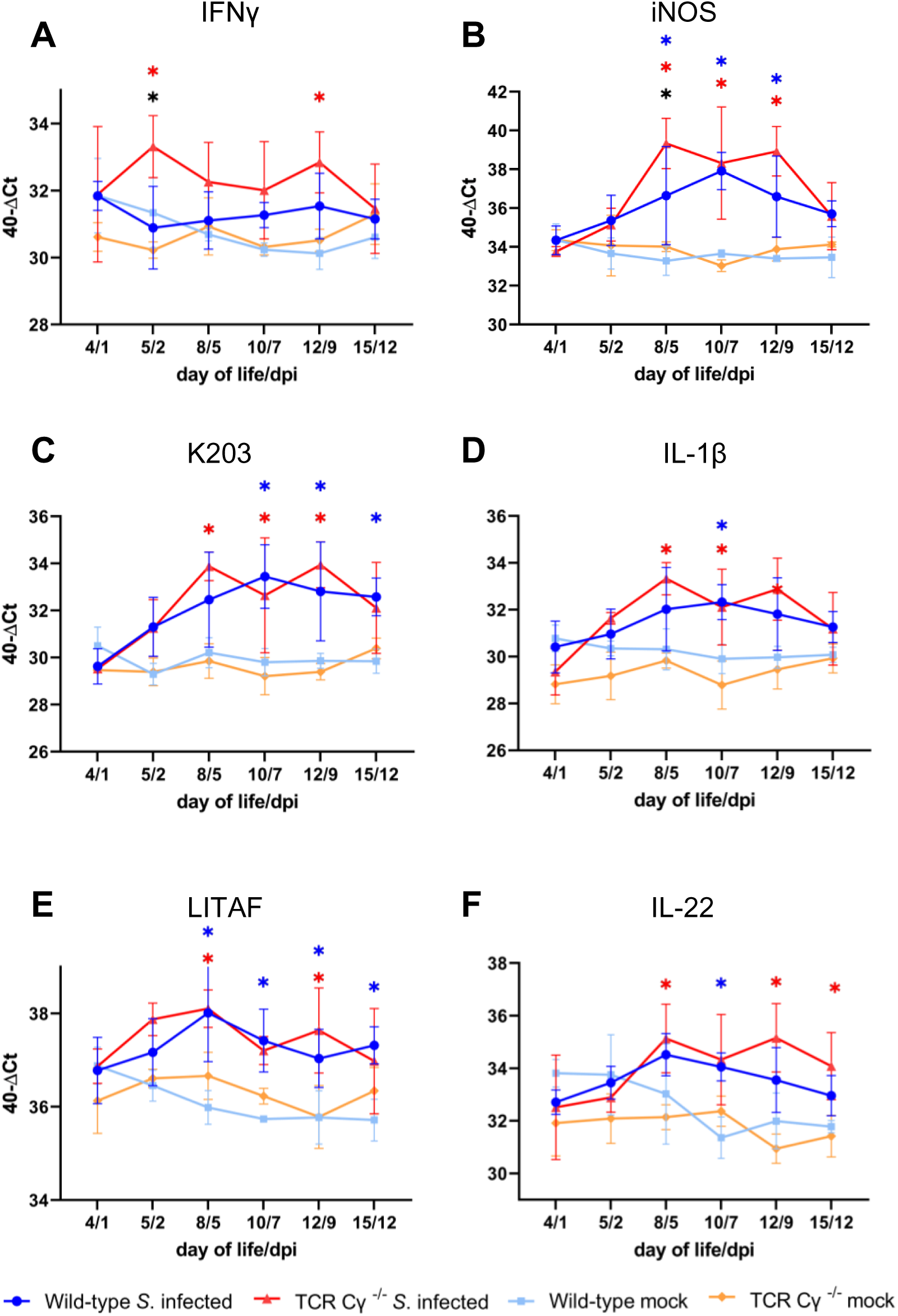
RT qPCR analysis of pro-inflammatory genes in cecal tissue of wild-type and TCR Cγ^−/−^ chickens following *Salmonella* Enteritidis infection. Transcription levels of IFN-γ, iNOS, K203, IL-1β, LITAF, and IL-22 (A-F) were measured in cecal tissues from wild-type and TCR Cγ^−/−^ chickens following *Salmonella* and mock infection. Data represent the mean 40-ΔCt ± standard deviation; n = 2-6. * indicates significant difference between *Salmonella*-infected wild-type and TCR Cγ^−/−^ chickens. */* indicate a significant difference between *Salmonella*-infected birds to their respective mock-infected controls, p < 0.05.

In contrast, anti-inflammatory cytokine transcription, such as TGFβ and IL10, remained largely unchanged following infection (Fig. 10). However, the immunomodulatory cytokine IL-21 showed increased expression in both chicken lines starting from 5 dpi. Analysis of cytotoxic effector molecules revealed significant differences in granzyme transcription levels between infected and control birds at 9 and 12 dpi. Transcription levels of perforin were also upregulated upon infection but with no significant differences between the infected and the control birds (Fig. 10).

**Figure 10:**
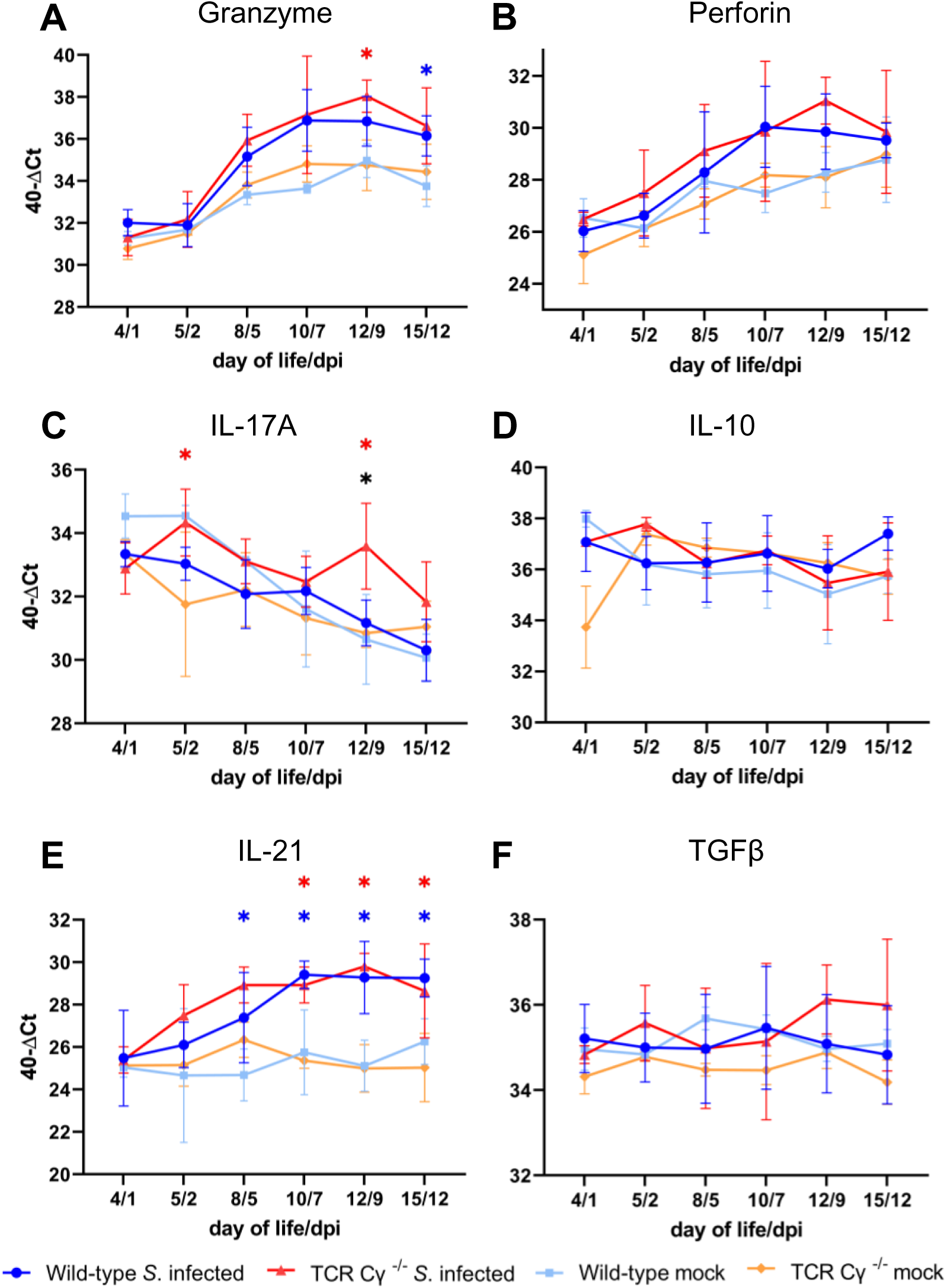
RT qPCR of immune related genes in cecal tissue of wild-type and TCR Cγ^−/−^ chickens following *Salmonella* Enteritidis infection. Transcription levels of granzyme, perforin, IL-17A, IL-10, IL-21, and TGFβ (A-F) were measured in cecal tissues from wild-type and TCR Cγ^−/−^ chickens following *Salmonella* and mock infection. Data represent the mean 40-ΔCt ± standard deviation; n = 2-6. * indicates a significant difference between *Salmonella*-infected wild-type and TCR Cγ^−/−^ chickens. */* indicate a significant difference between *Salmonella*-infected birds to their respective mock-infected controls, p < 0.05.

## 4. Discussion

Previous studies with *Salmonella*-infected wild-type chickens revealed an increased percentage of γδ T cells and therefore postulated a function of these enigmatic cells during the immune defense against *Salmonella* (38). However, the actual significance of γδ T cells in eliminating *Salmonella* from the avian host remained unclear. In this study, we demonstrate for the first time that γδ T cells can contribute to the protection of chickens against the *Salmonella* infection. Moreover, the CD8αα^++^ γδ T cell population, which has been observed to increase in wild-type chickens after *Salmonella* infection, appears to be compensated by a non-γδ T cell population, which is predominantly composed of αβ T cells in TCR Cγ^−/−^ chickens.

In our animal experiments, three-day-old γδ T cell knockout and wild-type chickens were infected with *Salmonella* Enteritidis. The infectious dose used resulted in a stable *Salmonella* colonization of the cecal lumen without significant differences between the chicken lines. Compared to previous studies, in which one day old wild-type chicks were infected with the same *Salmonella* strain and dose (9), we observed lower *Salmonella* colonization within the cecal lumen and reduced bacterial invasion into the liver of the chickens in this study. This disparity in results may be due to the age difference of the animals at the time of infection. The immune system and cecal microbiota of young chicks develop rapidly after hatching, and 3-day-old chickens are better protected against intestinal infections than newly hatched birds (4–6, 65).

Nonetheless, our results showed a significantly higher *Salmonella* invasion into the cecal wall and liver of *Salmonella*-infected TCR Cγ^−/−^ chickens than of wild-type animals, particularly at the onset of infection. This finding is consistent with studies in mice, which similarly reported increased translocation of *Salmonella* across the intestinal epithelium in the absence of γδ T cells during the early stages of infection (51, 66). The cfu in the liver of knockout animals even reached levels comparable to those observed in infected day-old wild-type chicks of previous studies (9). Thus, we show for the first time that γδ T cells have a decisive influence on the invasion and dissemination of *Salmonella* in young chickens. The γδ T cells may exert protective effects in multiple ways. Firstly, γδ T cells possess innate immune cell characteristics, can directly kill infected cells, recruit neutrophils and activate phagocytes (23, 67–69). Additionally, γδ T cells secrete a range of cytokines to regulate and shape the immune response (34, 70). Secondly, γδ T cells migrate from the thymus to the intestine during the embryonic development, starting at embryonic day 15. This is even significantly earlier than αβ T cell migration (71). Thus, γδ T cells seem to fill the immunological gap shortly after hatching, when the immune functions of αβ T cells are not yet available. Thirdly, the TCR Cγ^−/−^ chickens in this study may have had an impaired epithelial barrier integrity due to the absence of γδ T cells. In mouse and human intestinal tissue, γδ T cells represent an essential intraepithelial population, contributing to immune surveillance and maintenance of the intestinal integrity (72–74). In *Salmonella*-infected mice, γδ T cells maintain the integrity of the epithelial tight junctions, thereby limiting bacterial invasion into the gut mucosa (51, 66, 75, 76). However, whether and to what extent avian γδ T cells actually contribute to the maintenance of the intestinal barrier remains to be elucidated. In the absence of infection, Heyl et al. (40) reported no notable differences in intestinal morphology between TCR Cγ^−/−^ chickens and their wild-type counterparts.

In mice, other lymphocyte subsets can partially compensate for the absence of γδ T cells (49, 77, 78). For non-infected TCR Cγ^−/−^ chickens, a significant increase in the CD8αα^+^ αβ T cell population has been reported (40). In the present study, no significant differences were found in the number of CD8αα^+^CD4^-^ αβ T cells in the blood and spleen of healthy TCR Cγ^-/-^ and wild-type chickens. However, shortly after the increase of γδ T cells in wild-type chickens post *Salmonella* infection, a significantly enhanced number of previously undescribed non-γδ T cells was detected in blood and spleen of infected TCR Cγ^−/−^ chickens. These non-γδ T cells of γδ T cell knockout chickens had the same CD8αα^++^ sub-phenotype, together with an increased activation level (CD25^+^), as the elevated γδ T cells of wild-type birds. Based on these findings, we suggest that the non-γδ T cells may potentially substitute the functions of γδ T cells in the TCR Cγ^−/−^ chickens after *Salmonella* infection. This hypothesis is supported by the fact that the increase in non-γδ T cells in knockout animals coincided with a substantial reduction in *Salmonella* counts in the liver.

Previous research has reported that the γδ T cell population includes lymphocytes that share numerous phenotypic, functional, and homeostatic characteristics with their αβ T cell counterparts (79). To investigate whether αβ T cells potentially compensate for the absence of γδ T cells and their functions in the knockout animals after *Salmonella* infection, the emergence of non-γδ and αβ T cell subsets was compared. We found that at least parts of the CD8αα^++^CD4^-^ non-γδ T cells rising after infection may indeed be αβ T cells (CD4^-^) in *Salmonella*-infected TCR Cγ^−/−^ chickens. This suggests a compensatory role for CD8αα^++^CD4^-^ αβ T cells in the immune response against *Salmonella* in chickens lacking γδ T cells. However, the number of CD8αα^++^ αβ T cells detected did not completely reach the levels of CD8αα^++^CD4^-^ non-γδ T cells found in infected knockout animals.

The increased CD8αα^++^CD4^-^ non-γδ T cell population in *Salmonella*-infected TCR Cγ^−/−^ chickens may include additional, yet unidentified cell populations beyond Vβ1 and Vβ2 αβ T cells. Of relevance are natural killer cells, which express CD8 but lack T or B cell lineage-specific antigens (62, 80). NK cells comprise approximately 30% of intestinal intraepithelial lymphocytes and show enhanced activation following *Salmonella* infection (62, 81, 82). However, despite their prominence during embryonic development, NK cell abundance in blood and spleen of adult birds remains relatively low, representing only 0.5% to 1% of the lymphocyte population (62, 80). Our study identified a low abundance population of CD8α^+^ lymphocytes lacking TCR1, TCR2, TCR3, Bu1 and K1 expression (termed NK-like cells herein) in blood and spleen. Consistent with existing literature (62, 80), a higher percentage of these cells was found in the cecum. While the overall NK-like cell population remained comparable between chicken lines, we observed significantly more CD8^+^ NK-like cells expressing the CD25 antigen in the blood of infected knockout animals compared to wild-type animals, with a similar trend in the cecum and spleen. The expression of the CD25 antigen (IL-2Rα) by human NK cells has been correlated with induced cytotoxic activity and cytokine secretion (83). Thus, NK-like cells appear to contribute to the functional compensation of γδ T cells in knockout chicken post infection. Indeed, Göbel et al. (62) previously postulated that γδ T cells and NK cells have redundant functions. However, whether the detected NK-like cells belong to the described CD8αα^++^CD4^-^ non-γδ T cell population or may represent an additional immune cell subset contributing to the compensation of missing γδ T cells needs to be clarified in further studies. Notably, NK cells do not consistently express the CD8 antigen on their surface, but also other markers, including CD56, CD16, and 20E5 (82, 84, 85). Further research is required to conclusively determine the compensatory capacity of NK cells in *Salmonella*-infected TCR Cγ^−/−^ chickens.

Results from this study further indicate that γδ T cell compensation in knockout chickens is not restricted to T or NK-like cells. There was a trend towards increased numbers of blood monocytes in TCR Cγ^−/−^ chickens compared to wild-type chickens after infection. This observation suggests enhanced cellular migration to sites of inflammation in infected knockout chickens. In tissues, monocytes differentiate into macrophages, which can typically be activated by γδ T cells during immune responses to infections (67). The elevated number of circulating monocytes may thus contribute to the attempt to compensate for the absence of γδ T cells and the resulting deficit of activated resident macrophages in tissues of knockout birds.

Regulation or dysregulation of specific immune mediators are also indicators of compensatory measures taken by an immunologically impaired and infected host. Analysis of immune gene transcription in the cecum showed elevated expression of IFN-γ and iNOS in TCR Cγ^−/−^ chickens compared to wild-type chickens post infection. Previous studies have demonstrated that the transcription levels of proinflammatory cytokines, including IFN-γ and iNOS, depend on the invasiveness of the *Salmonella* strain into the tissues of infected chickens (9, 63). The findings of our study support this correlation, as higher *Salmonella* counts were also found in the tissues of TCR Cγ^−/−^ chickens compared to wild-type animals. Notably, the transcription rates of other proinflammatory cytokines, such as LITAF and IL-1β, did not differ significantly between the infected TCR Cγ^−/−^ chickens and wild-type animals. This indicates that the knockout animals are only able to compensate for the loss of γδ T cells at certain regulatory levels after infection.

To determine whether the newly described CD8αα^++^CD4^-^ non-γδ T cells are able to replace the functions typically performed by γδ T cells distinct T cell populations were sorted from PBMCs and the transcription of key immune genes was analyzed. The results suggest that CD8αα^++^CD4^-^ non-γδ T cells are not only functionally equivalent to their CD8αα^++^ γδ T cell counterparts, but may surpass them in responsiveness following infection. Specifically, CD8αα^++^CD4^-^ non-γδ T cells of TCR Cγ^−/−^ chickens showed higher transcription rates of IL-17A, IL-2Rα and IFN-γ compared to CD8αα^++^ γδ T cells of wild-type chickens during the early stages of infection. Over time, CD8αα^++^ γδ T cells increased their transcription levels, eventually reaching rates comparable to their non-γδ T cell counterparts. This response dynamic of avian CD8αα^++^ γδ T cells aligns with previous studies showing an upregulation of IFN-γ in response to *Salmonella* infection (34, 38).

IL-17A is essential for the recruitment and activation of macrophages and neutrophils as first line of defense against salmonellosis (86). Additionally, IL-17A promotes pro-inflammatory cytokines at mucosal barriers, thereby enhancing pathogen control (87, 88). Notably, γδ T cells are a major source of IL-17A in the early stages of inflammatory and infectious responses. During *Listeria monocytogenes* infection in mice, Hamada et al. (42) showed an elevated absolute number of IL17A producing γδ T cells at 5 dpi. In chickens, Walliser and Göbel (36, 89) first described IL-17 producing CD4^+^ αβ T helper_17_ and γδ T cells. Our findings confirmed that avian CD8αα^++^ γδ T cells not only increase in number but also upregulate IL-17A transcription, highlighting an essential regulatory role during infection. Interestingly, our study revealed an increased IL-17A transcription by CD8αα^++^ γδ T cells particularly at later time points post infection, when innate immune defenses to control bacterial invasion were already advanced. These observations show a contribution of γδ T cells to IL-17A production beyond early innate immunity, but also during later adaptive immune responses to *S*. Enteritidis infection (90).

CD8αα^++^CD4^-^ non-γδ T cells in chickens not only produce IL-17A, but also respond more rapidly than γδ T cells during *Salmonella* infection. Previous studies have reported on the role of CD4^-^ non-γδ T cells as additional IL-17A-producers during *Salmonella* infection (90). While CD8αα^++^CD4^-^ non-γδ T cells are present in low numbers in wild-type chickens, in our study they may compensate for the loss of γδ T cell-derived IL-17A in TCR Cγ^−/−^ chickens through both numerical expansion and early activation post infection. The high transcription rate of IL-2Rα in CD8αα^++^CD4^-^ non-γδ T cells, indicating enhanced activation and proliferation, further supports the crucial role of these cells for immune regulation in the absence of γδ T cells (91, 92).

In conclusion, the present study emphasizes the pivotal role of γδ T cells in mediating the early immune response of chickens to *Salmonella* Enteritidis infection. Other cell populations, particularly CD8αα^++^CD4^-^ non-γδ T cells, but also NK-like cells, appear to be utilized by the host to compensate for the absence of γδ T cells. Especially, the CD8αα^++^CD4^-^ non-γδ T cells, consisting at least in part of αβ T cells, represent a unique and pre-activated cell population with high potential for proliferation and protective immune response in *Salmonella*-infected TCR Cγ^−/−^ chickens. Beyond numerical replacement of the missing γδ T cells, compensatory efforts by the host also make an impact on the functional and regulatory level of the immune response. Nevertheless, γδ T cell knockout chickens exhibit impaired early immune protection, resulting in comparatively stronger *Salmonella* dissemination throughout the body. Further research is needed to assess whether the identified compensatory elements are only indicative of a dysregulated, functionally impaired immune response, or whether they are able to effectively replace the γδ T cells in conferring a protective host response against *Salmonella*. Understanding these dynamics could be deployed to enhance the resilience of chickens to infections across various tissues and stages of infection.

## Data availability statement

The data presented in the study are publicly available and can be found here:

## Ethics statement

The animal study was approved by the Ethics Committee for Animal Experiments and Animal Welfare of the State of Thuringia, Germany, identification code: BFI-20-001. The study was conducted in accordance with the local legislation and institutional requirements.

## Author contributions

FT: Investigation, Formal analysis, Conceptualization, Data curation, Visualization, Writing – original draft, Writing – review and editing. AB: Supervision, Conceptualization, Funding acquisition, Methodology, Investigation, Project administration, Writing – original draft, Writing – review and editing. UM: Methodology, Investigation, Writing – review and editing. TvH: Resources, Writing – review and editing. BS: Resources, Writing – review and editing. CM: Conceptualization, Resources, Writing – review and editing.

## Acknowledgments

We thank Katrin Schlehahn for excellent technical assistance especially with the flow cytometry and RT-PCR experiments. Special thanks go to the staff of the animal production and experimentation unit at the ‘Friedrich-Loeffler-Institut’ Jena. Also, we thank Marie-Sophie Sädler, Norman Müller and Nadine Taupitz for their experimental assistance during this study.

## Publisher’s note

All claims expressed in this article are solely those of the authors and do not necessarily represent those of their affiliated organizations, or those of the publisher, the editors and the reviewers. Any product that may be evaluated in this article, or claim that may be made by its manufacturer, is not guaranteed or endorsed by the publisher.

## Supplementary material

The Supplementary Material for this article can be found online at:

