## Supplemental Figures for "Compensatory Mechanisms in γδ T Cell-Deficient Chickens Following *Salmonella* infection"

### Supplementary Material

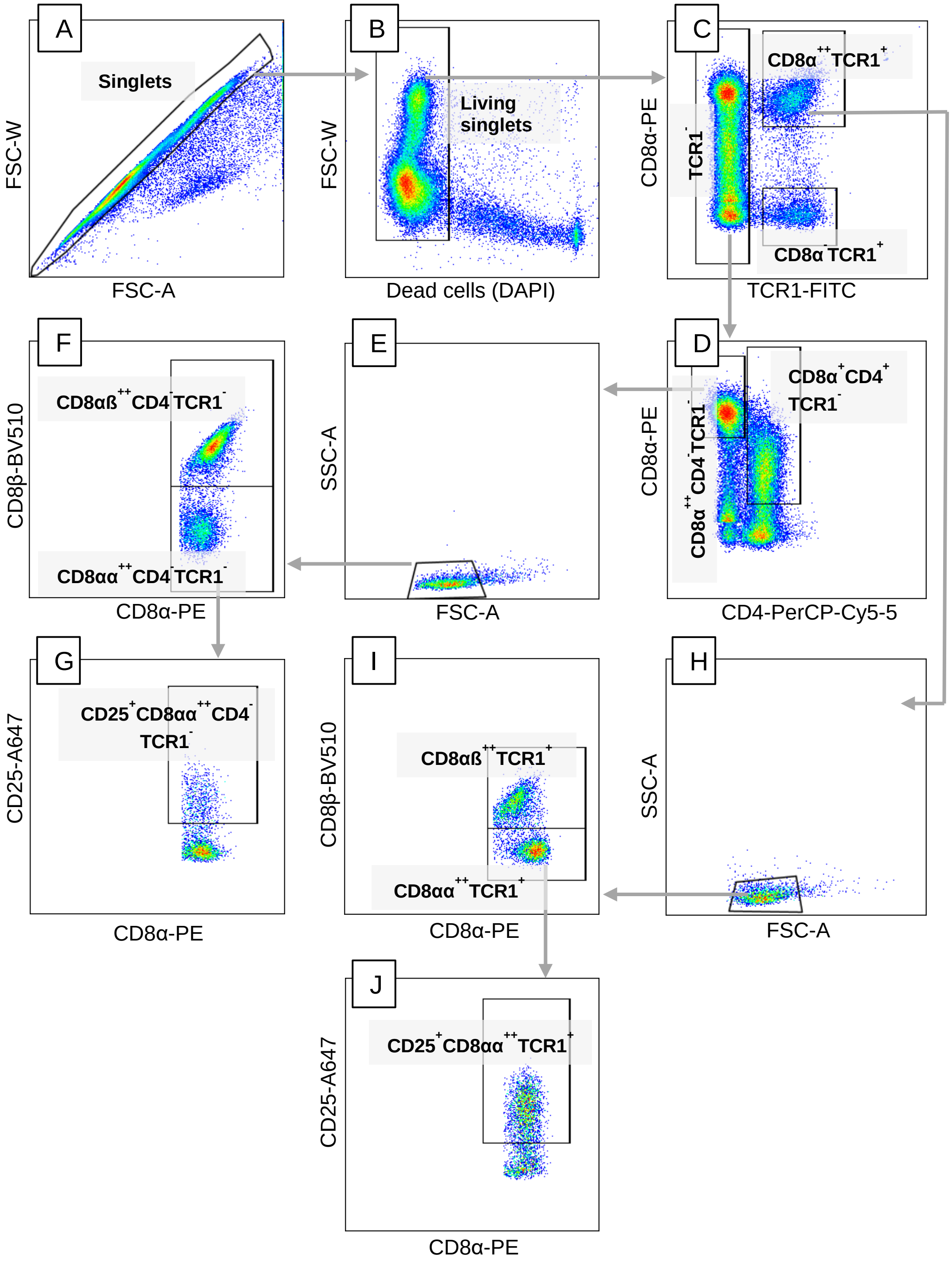


**Supplementary Figure 1. Representative gating strategy for identifying TCR1$^{+}$ and TCR1$^{-}$ T cell subsets.** Following the exclusion of doublets and dead cells (A, B), γδ T cells (TCR1$^{+}$) and non-γδ T cells (TCR1$^{-}$) were gated and distinguished based on CD8α and TCR1 expression (C). TCR1$^{-}$ cells were further subdivided into CD4-positive and CD4-negative subsets (D). All lymphocyte subpopulations were validated by back-gating using an SSC/FSC dot plot (E, H). CD8α$^{++}$TCR1$^{+}$ and

$CD8\alpha^{++}CD4^{-}TCR1^{-}$ T cells were further classified into CD8αα-positive and CD8αß-positive subsets (F, I). The activation status of these subsets was assessed by CD25 expression (G, J).

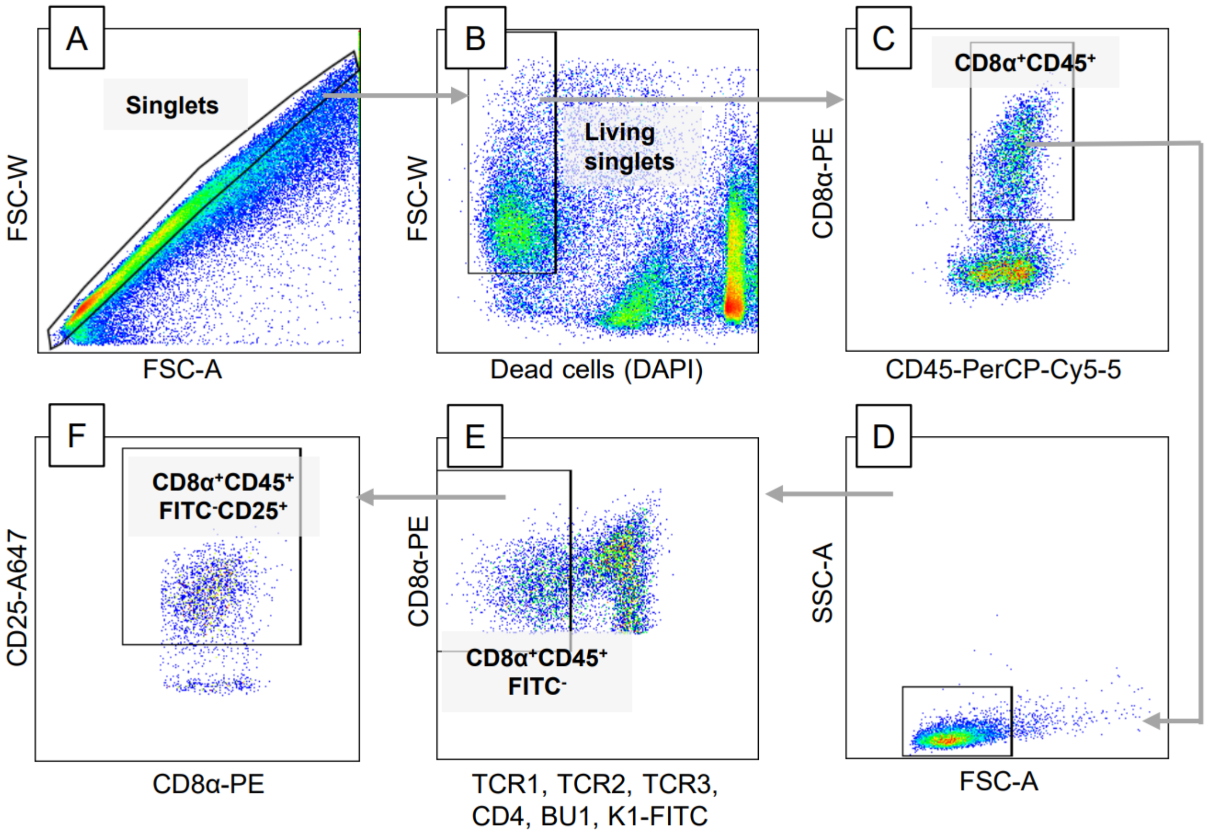


**Supplementary Figure 2. Representative gating strategy for identifying NK-like lymphocyte subsets.** Doublets and dead cells were excluded from the analysis (A, B). CD8α$^+$ leucocytes (CD45$^+$) were gated based on the dot-plot showing CD8α and CD45 expression (C). The lymphocyte subpopulation was confirmed by back-gating using an SSC/FSC dot plot (D). Cells expressing T cell, B cell, and monocyte lineage markers (FITC-conjugated) were excluded, identifying the CD8α$^+$CD45$^+$FITC$^-$ subset (E). The activation status of the CD8α$^+$CD45$^+$FITC$^-$ cell subset was assessed by CD25 expression, visualized on a CD25 versus CD8α plot (F).

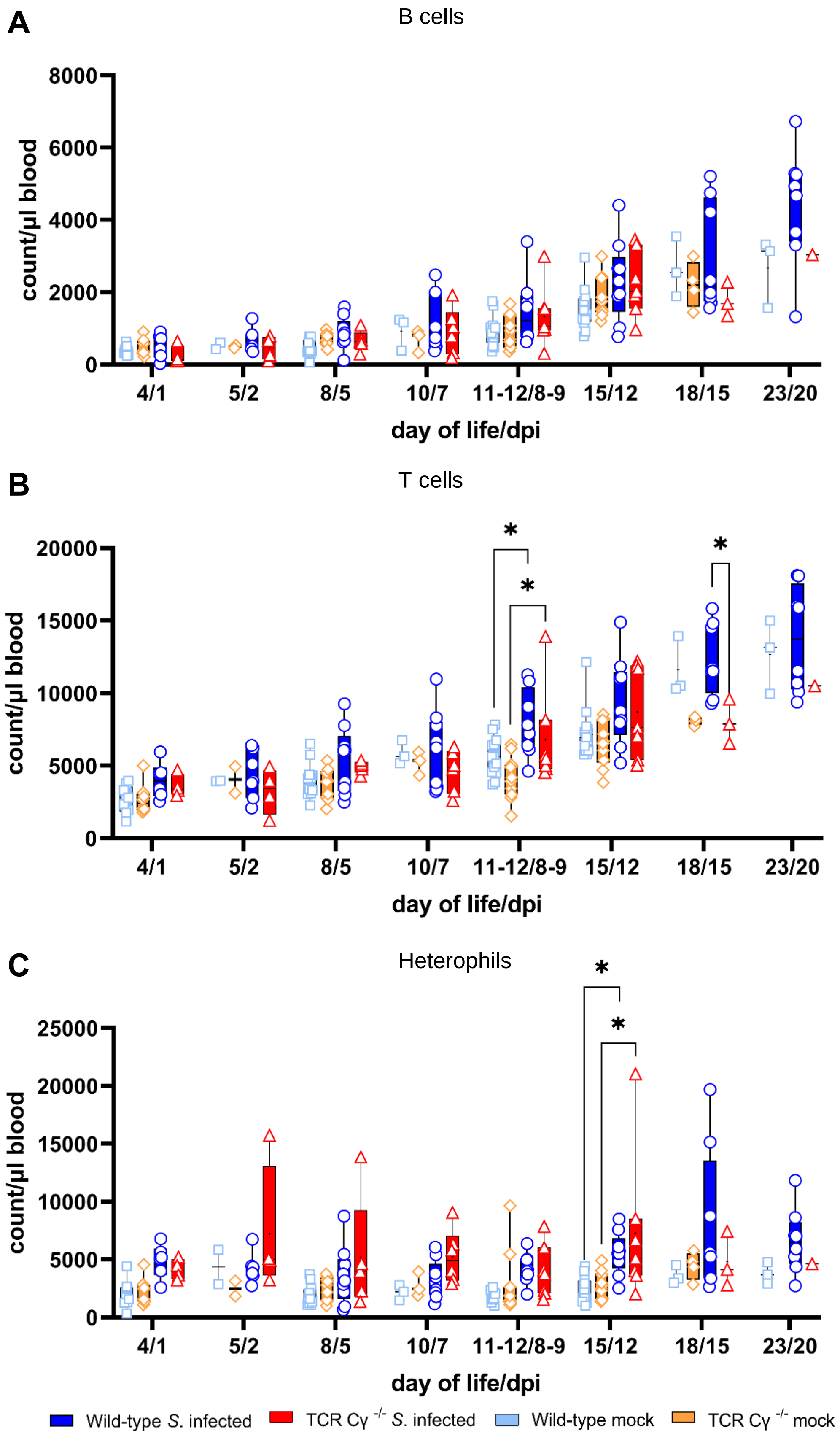
A
B cells
count/µl blood
8000
6000
4000
2000
0
4/1
5/2
8/5
10/7
11-12/8-9
15/12
18/15
23/20
day of life/dpi
B
T cells
count/µl blood
20000
15000
10000
5000
0
*
*
*
4/1
5/2
8/5
10/7
11-12/8-9
15/12
18/15
23/20
day of life/dpi
C
Heterophils
count/µl blood
25000
20000
15000
10000
5000
0
*
*
4/1
5/2
8/5
10/7
11-12/8-9
15/12
18/15
23/20
day of life/dpi
Wild-type S. infected
TCR Cγ -/- S. infected
Wild-type mock
TCR Cγ -/- mock

**Supplementary Figure 3. Flow cytometric analysis of leukocytes in whole blood following *Salmonella* Enteritidis infection.** Absolute numbers of viable B cells (A), T cells (B) and heterophils (C) were measured over time in blood from wild-type and TCR $C\gamma^{-/-}$ chickens following *Salmonella* and mock infection. Data are presented as minimum and maximum cell counts, with median indicated; n = 1-13. * indicates significant differences between chicken groups, $p < 0.05$. No data available for mock-infected TCR $C\gamma^{-/-}$ chickens at 20 dpi.

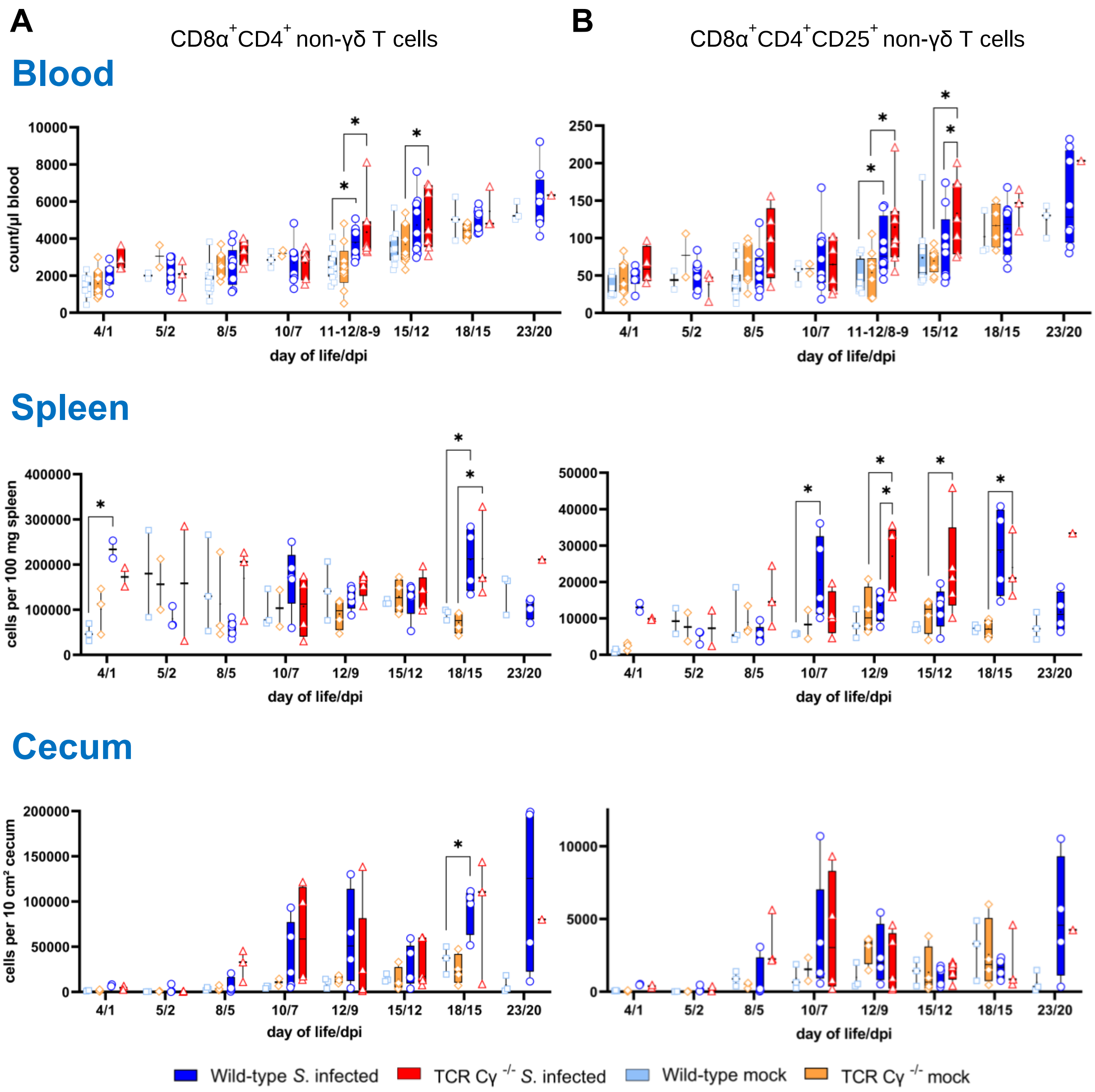


**Supplementary Figure 4. Flow cytometric analysis of CD8α$^{+}$CD4$^{+}$ T cells in *Salmonella* Enteritidis infected chickens.** Absolute numbers of CD8α$^{+}$CD4$^{+}$ (A) and CD8α$^{+}$CD4$^{+}$CD25$^{+}$ (B) non-γδ T cells are shown for blood (n = 1-13), spleen (n = 1-5), and cecum (n = 1-5) from wild-type and TCR Cγ$^{-/-}$ chickens following *Salmonella* and mock infection. Data are presented as minimum and maximum cell counts, with median indicated. * indicates significant differences between chicken groups, $p < 0.05$. No data available for mock-infected TCR Cγ$^{-/-}$ chicken at 20 dpi.
